# Lamin A/C regulates the compartment-specific contributions of immune and stromal cells to intestinal inflammation and colitis-associated colon cancer

**DOI:** 10.64898/2026.07.03.735779

**Authors:** Raquel Gómez-Bris, Marina Ortega-Zapero, Beatriz Herrero-Fernández, Víctor Fanjul, Nicolás de la Madrid de Vega, Sara Morán de Bustos, Irene Moreno-Aperribay, Virginia Zorita, Héctor Sánchez-Martinez, Lauri Polari, Alicia Usategui, Marta Amorós-Pérez, Pilar Gonzalo, Markku Voutilainen, Markku Kallajoki, Jesús Vázquez, Juan Antonio López, Jose Luis Pablos, Gabriel Criado, Silvia M Arribas, Carlos Silvestre-Roig, Francisco Sánchez-Madrid, Vicente Andrés, Diana M Toivola, Angela Saez, José M. González-Granado

**Author notes:** Authors for correspondence. AS; JMGG. Equally contribution (Raquel Gómez-Bris, Marina Ortega-Zapero).

## Abstract

Inflammatory bowel disease (IBD) arises from dysregulated crosstalk between innate immune, adaptive immune, and stromal compartments, yet the compartment-specific mechanisms driving tissue injury and tumorigenesis remain incompletely defined. To address this gap, we used conditional knockout and overexpression mouse models together with human IBD biopsy analysis to dissect the compartment-specific roles of lamin A/C in intestinal inflammation and colitis-associated tumorigenesis. Pan-hematopoietic lamin A/C deletion attenuated acute dextran sulfate sodium (DSS)-induced colitis. Myeloid-specific lamin A/C deletion ameliorated chronic colitis and was associated with altered dendritic cell (DC) programs, enhanced regulatory T cell (Treg) responses, and reduced effector T cell activation. Adoptive transfer of lamin A/C-deficient bone marrow-derived DCs recapitulated this reduced-damage phenotype in DSS colitis, while proteomic profiling revealed reduced antigen-processing and inflammatory programs together with enhanced metabolic and mucosal defense pathways. T cell-specific lamin A/C deletion reduced the Th1/Treg ratio and limited tumor development by suppressing chronic inflammation, whereas T cell-specific lamin A/C overexpression promoted severe Th1-skewed pathology, sustained intestinal inflammation, and increased colitis-associated tumor burden. Stromal fibroblast-specific lamin A/C deletion generated a tissue-protective niche characterized by enhanced epithelial barrier gene expression, regulatory cytokine production, and remodeling of the local immune milieu. Human IBD biopsies revealed compartment-specific lamin A/C alterations consistent with the murine findings. In lamina propria CD3⁺ T cells, lamin A/C levels were blunted in IBD and associated with local histological severity rather than IBD diagnosis, whereas epithelial lamin A/C showed a steeper crypt-axis spatial gradient in a Crohn’s disease-specific pattern. Together, these findings identify lamin A/C as a cell-type- and context-dependent regulator of intestinal inflammation and tumorigenesis.

## Introduction

Inflammatory Bowel Disease (IBD), encompassing Crohn’s disease (CD) [1] and ulcerative colitis UC [2], is a chronic, relapsing inflammatory disorder of the gastrointestinal tract [3]. Its pathogenesis involves a dysregulated interplay between host immunity, the intestinal microbiota, and the mucosal barrier. Innate immune cells such as neutrophils, dendritic cells (DCs), and macrophages initiate inflammation [4, 5, 6, 7], whereas adaptive CD4^+^ T cells, particularly Th1 and Th17 subsets, sustain chronic tissue damage [8, 9, 10]. Stromal fibroblasts and the epithelium further contribute through cytokine and chemokines production and matrix remodeling [11, 12, 13, 14, 15]. Despite this understanding, how each cellular compartment differentially contributes to IBD initiation, progression, and heterogeneity remains incompletely defined, hindering the development of targeted therapies. Experimental models enabling cell-type-specific interrogation are crucial to clarify this compartmental organization [12, 16, 17].

The nuclear lamina component lamin A/C, encoded by *LMNA*, represents a promising molecular target for IBD. Its expression is largely absent in naïve lymphocytes but transiently induced upon activation and dynamically regulated in the myeloid compartment by maturation state and local microenvironment. Its expression is largely absent in naïve lymphocytes but transiently induced upon activation. In myeloid cells, lamin A/C abundance is dynamically remodeled during differentiation and activation, often increasing during monocyte-to-macrophage maturation but decreasing upon pro-inflammatory macrophage activation; granulocyte-macrophage colony-stimulating factor (GM-CSF)–derived DCs can also express high lamin A/C. In both lineages, its levels are microenvironment-dependent [18, 19, 20, 21, 22]. Beyond its structural role, lamin A/C influences nuclear mechanics, chromatin organization, and signaling pathways that shape cellular responses [23, 24]. In T cells, it affects immunological synapse formation [19, 25] and differentiation programs [26, 27], while in DCs it modulates nuclear deformability and mechanics [28] and inflammatory responses [20, 29]. These observations suggest lamin A/C may regulate immune cell function in a cell-specific and context-dependent manner. Based on this rationale, we hypothesized that lamin A/C differentially modulates the pathogenic or tolerogenic potential of distinct cellular lineages in intestinal inflammation. To test this, we employed a panel of conditional knockout and overexpression mouse models targeting pan-hematopoietic, myeloid, T cell, and stromal fibroblast populations, together with human IBD biopsy analysis, to dissect the compartment-specific roles of lamin A/C in intestinal inflammation and colitis-associated cancer.

## Results

### 1. Increasing pan-hematopoietic lamin A/C levels promotes susceptibility to acute DSS-induced colitis

To investigate whether lamin A/C levels within the immune compartment modulate intestinal inflammation, acute colitis was induced by a single DSS cycle in mice with pan-hematopoietic deletion (Vav1-KO) or overexpression (Vav1-OE) of *Lmna* (Suppl. Fig. 1A). Vav1-KO mice developed a significantly attenuated disease course compared with wild-type (WT) littermates, as reflected by reduced weight loss and lower disease activity index (DAI) throughout DSS exposure (Suppl. Fig. 1B, C). In contrast, Vav1-OE mice exhibited exacerbated colitis, characterized by increased weight loss and higher DAI relative to WT animals (Suppl. Fig. 1D, E). Collectively, these data demonstrate that lamin A/C dosage within the hematopoietic compartment quantitatively regulates susceptibility to acute DSS-induced intestinal inflammation.

### 2. Lamin A/C–dependent protection is preserved in the absence of adaptive lymphocytes

To determine whether the protective phenotype observed in Vav1-KO mice depended exclusively on adaptive lymphocytes, acute DSS colitis was induced in both Vav1-KO-Rag1 mice and Vav1-WT-Rag1 controls. Vav1-KO-Rag1 mice lack mature T and B cells while also lacking lamin A/C in the remaining hematopoietic lineages, whereas Vav1-WT-Rag1 controls maintain physiological lamin A/C levels across all immune cells but also lack adaptive lymphocytes due to Rag1 deficiency (Suppl. Fig. 1F). Following DSS exposure, Vav1-KO-Rag1 mice exhibited reduced weight loss and lower DAI compared with Vav1-WT-Rag1 controls (Suppl. Fig. 1G, H). These findings indicate that lamin A/C deletion in non-lymphoid hematopoietic populations is sufficient to confer protection, supporting a broad immune-intrinsic role for lamin A/C in amplifying intestinal inflammation.

### 3. Myeloid-specific lamin A/C deletion attenuates DSS-induced colitis and modulates adaptive immune responses

Given the contribution of non-lymphoid immune cells, we next examined the role of lamin A/C in the myeloid compartment using LysM-KO mice which lack lamin A/C in myeloid cells. Colitis was induced using a single DSS cycle or a two-cycle DSS protocol to model acute and chronic disease, respectively (Fig. 1A). After a single DSS cycle, LysM-KO mice showed a milder disease course, with reduced body weight loss, lower DAI, and diminished colon shortening compared with controls (Fig. 1B–D). Histological analysis confirmed significantly reduced histological damage score in LysM-KO mice (Fig. 1E). In the chronic two-cycle DSS model, body weight trajectories were comparable between genotypes (Fig. 1B), but clinical DAI diverged during the second cycle, with LysM-KO mice accumulating lower clinical scores (Fig. 1C). Moreover, colon shortening remained reduced (Fig. 1F), and histological damage was significantly lower in LysM-KO (Fig. 1G).

**Figure 1.**
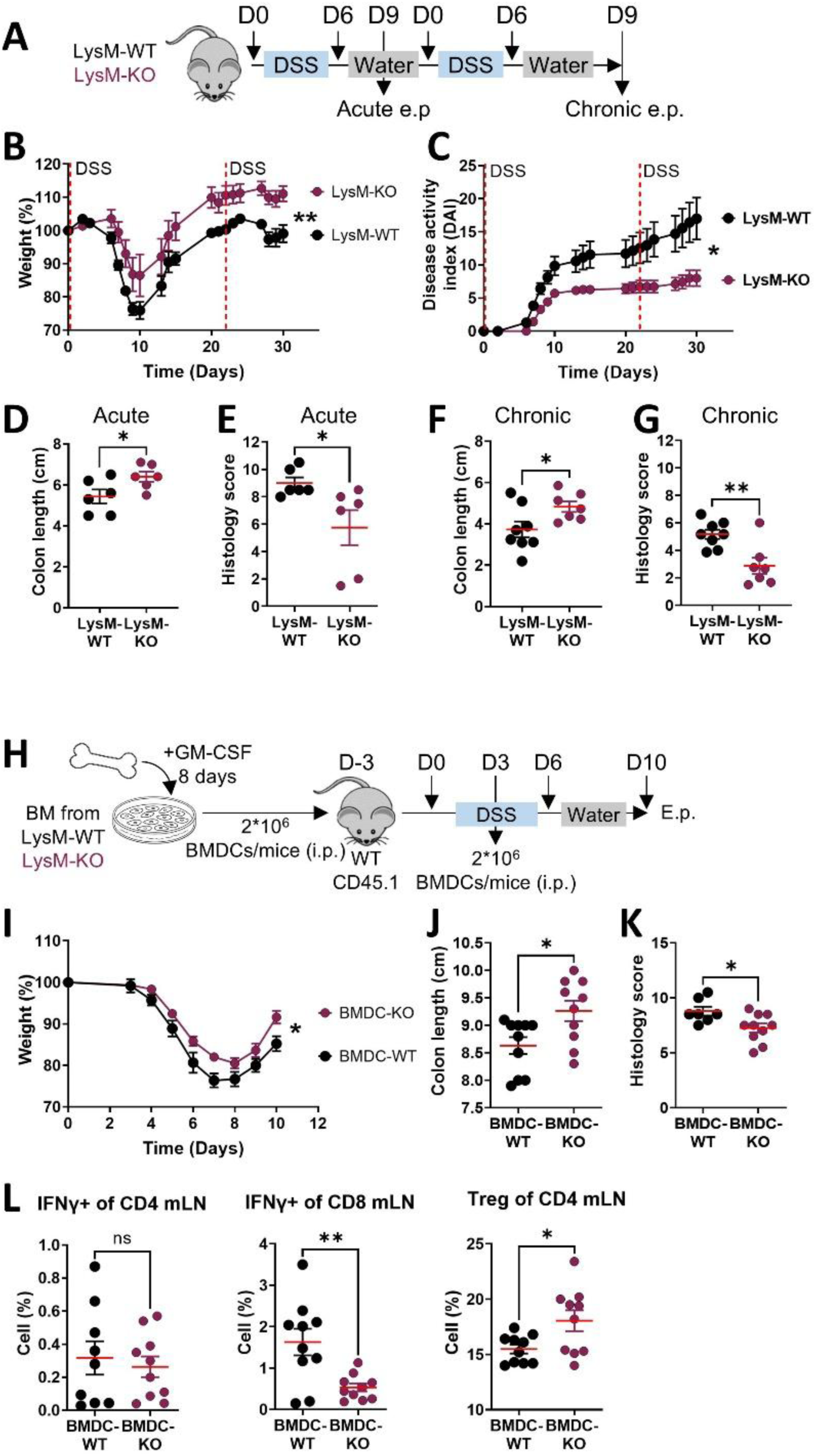
Myeloid-specific lamin A/C deletion attenuates acute and chronic DSS-induced colitis and lamin A/C–deficient bone marrow-derived dendritic cells (BMDCs) attenuate DSS-induced colitis and indirectly modulate adaptive T-cell responses. (A) Schematic representation of the DSS-induced colitis protocol used to evaluate the impact of myeloid lamin A/C deletion in wild-type (LysM-WT) and myeloid-specific lamin A/C–deficient (LysM-KO) mice using one-cycle acute DSS colitis or two-cycle chronic DSS colitis. Discontinuous vertical lines indicate the start of each DSS cycle. (B) Body weight evolution during the acute and chronic DSS cycles, expressed as percentage of day 0. (C) Disease activity index (DAI) during acute and chronic colitis. (D) Colon length measured at sacrifice after acute DSS cycle. (E) Blinded histopathological scores after acute DSS cycle. (F) Colon length measured at sacrifice after chronic DSS cycle. (G) Blinded histopathological scores after chronic DSS cycle. (H) Schematic representation of the adoptive transfer experimental design. BMDCs were generated in vitro with GM-CSF from WT and LysM-KO donor mice and transferred i.p. into WT CD45.1 recipient mice 3 days before DSS exposure and again on day 3 of DSS exposure; mice were sacrificed 10 days after DSS initiation. (I) Body weight evolution during DSS exposure, expressed as percentage of day 0. (J) Colon length measured at sacrifice. (K) Blinded histopathological scores at sacrifice. (L) Flow cytometric analysis of mesenteric lymph nodes (mLN) showing the frequency of IFNγ-producing CD4^+^ T cells, IFNγ-producing CD8^+^ T cells, and Foxp3^+^ regulatory T cells (Treg) among CD4^+^ T cells. Data are shown as mean ± SEM (n = 6–8 mice per genotype pooled from two independent experiments for WT/LysM-KO comparison; n = 5–7 mice per group; one representative experiment out of at least three independent experiments for BMDC adoptive transfer experiment). Statistical significance was assessed using two-way ANOVA with Tukey’s multiple-comparison test for longitudinal measurements and unpaired Student’s t-test for endpoint analyses; *p < 0.05, **p < 0.01, ns not significant. Endpoint: e.p. Intraperitoneal: i.p.

Given that LysM-driven deletion targets multiple myeloid subsets, we next assessed whether DCs alone were sufficient to confer protection using adoptive transfer (Fig. 1H). Adoptive transfer of lamin A/C-deficient bone marrow–derived DCs (BMDCs KO) to WT mice recapitulated the protective phenotype of LysM-KO mice, with reduced weight loss, attenuated colon shortening, and diminished histological damage score compared with WT recipients of BMDCs WT (Fig. 1I-K). At the cellular level, adoptively transferred BMDCs KO were associated with increased Treg frequencies and reduced IFNγ-producing CD8⁺ T cells (Fig. 1L), indicating indirect modulation of adaptive immunity.

Given that transferred GM-CSF–derived BMDCs are expected to encounter inflammatory and bacterial signals in the DSS-injured intestine, we next examined the impact of lamin A/C loss in lipopolysaccharide (LPS)-matured BMDCs. To this end, TMT-based quantitative proteomics was applied to LPS-matured WT and LysM-KO BMDCs. A total of 8,218 proteins were detected and quantified at false discovery rate (FDR) < 1%, and 8,035 remained after excluding artifacts, including contaminants and inconsistently expressed proteins in controls. Of these, 5,793 proteins were quantified with more than one unique peptide. After verifying correct sample normalization and group separation, a significance threshold for altered expression was set at z = ±1.2 (Supp. Fig. 2A-C). A total of 4.4% of proteins showed differential expression between KO and WT BMDCs (Supp. Fig. 3; Supp. Table 1). GSEA using the KEGG database revealed that KO BMDCs exhibited downregulation of pathways related to antigen engulfment, digestion, processing, and presentation, as well as signaling, detoxification, Th1/2-cell activation, and inflammatory disease. Conversely, these cells showed upregulated lipid, carbohydrate, and amino acid metabolism pathways, together with pathways related to mucosal defense and signaling (Fig. 2A, B). These results suggest that lamin A/C deficiency reprograms BMDCs toward a less inflammatory, more tolerogenic state, together with metabolic rewiring.

**Figure 2.**
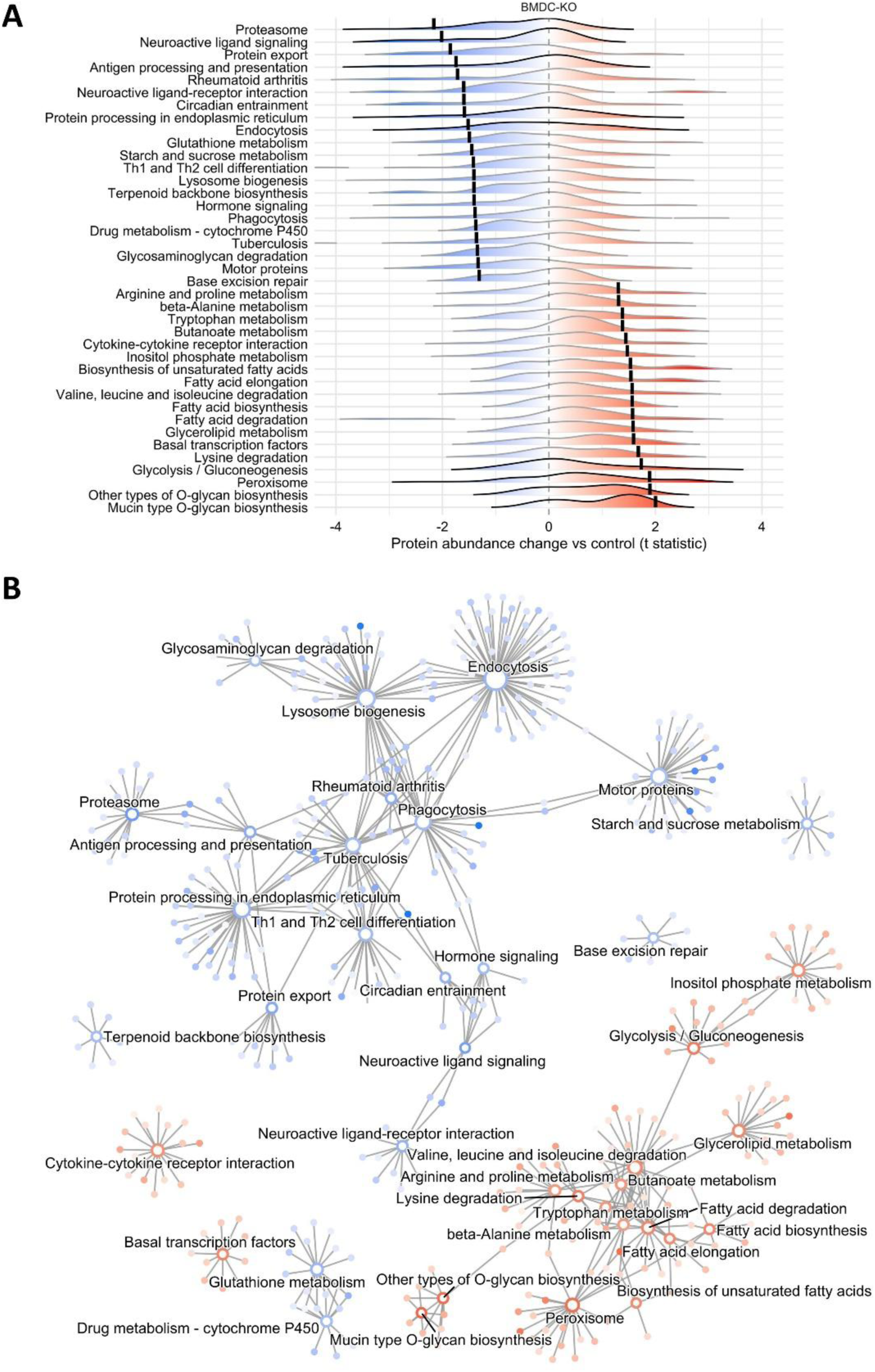
Quantitative proteomics examined in GM-CSF-generated-LPS-matured–lamin A/C–deficient bone marrow-derived dendritic cells (BMDC) suggest that lamin A/C deficiency induces a tolerogenic phenotype. Enriched KEGG pathways. A) Pathway ridgeline plot. Each ridge illustrates the density distribution of individual proteins within a significantly enriched category. The x-axis represents the relative protein abundance change (t-statistic) of BMDC-KO compared to WT; a shift to the right indicates upregulation, while a shift to the left indicates downregulation. The overall pathway-level shift is represented by the Normalized Enrichment Score (NES), marked as a vertical line on each ridge. |NES| > 1.3 was used to determine statistical differences. Categories meeting the strict significance threshold (adjusted p value < 0.05) are outlined in black. B) Concept network per group. Individual proteins are represented by color-filled nodes (t-statistic), whereas categories are represented by color-bordered nodes (NES). The internal gradient color reflects the individual protein t-statistics, with color intensity saturating at |t| or |NES| ≥ 3.

**Figure 3.**
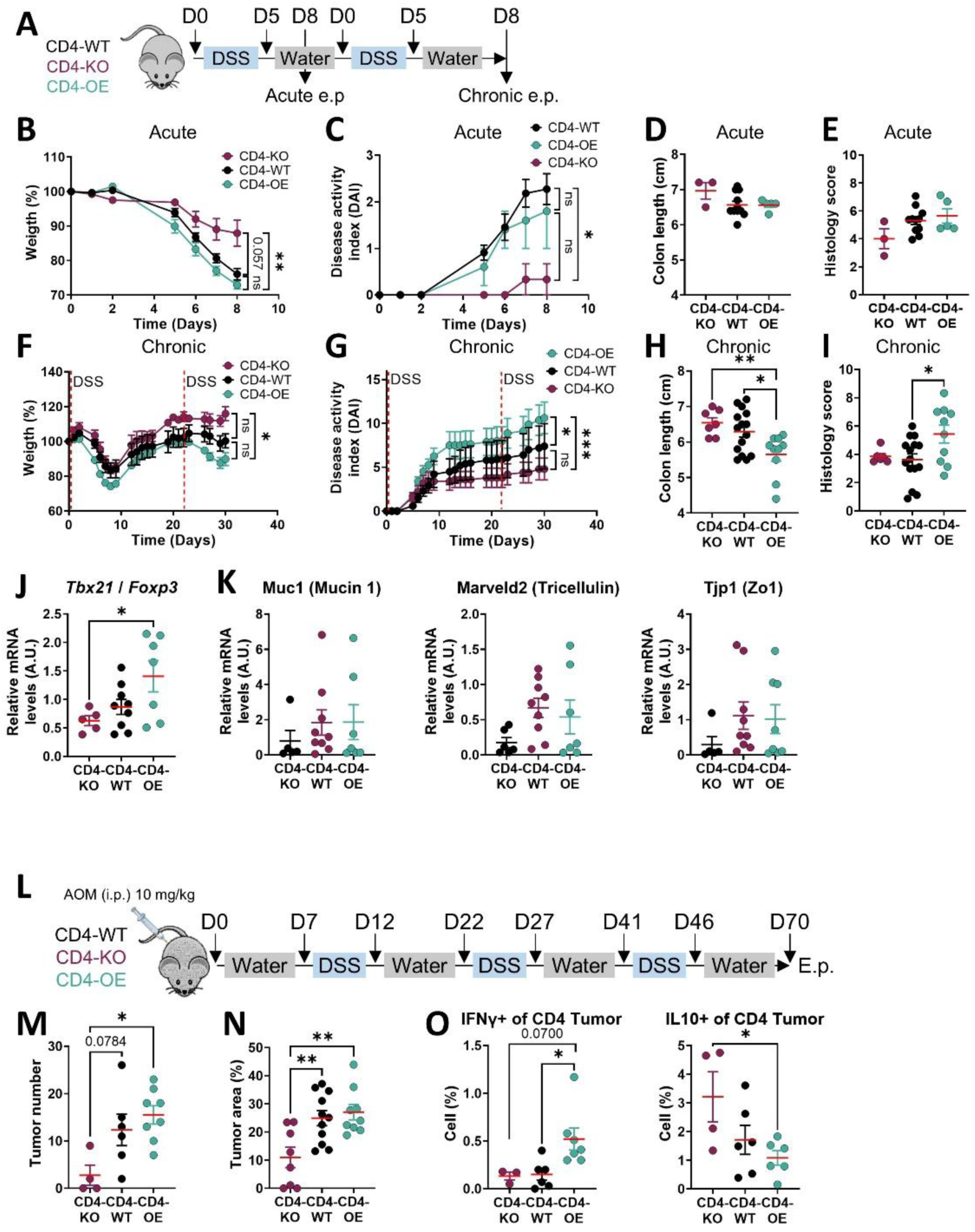
T cell–specific lamin A/C regulates intestinal inflammation and colitis-associated colorectal cancer development. (A) Schematic representation of the DSS-induced colitis protocols used to assess the role of T cell–intrinsic lamin A/C in wild-type (CD4-WT), T cell–specific lamin A/C–deficient (CD4-KO), and T cell–specific lamin A/C–overexpressing (CD4-OE) mice, using one-cycle acute DSS colitis or two-cycle chronic DSS colitis. Discontinuous vertical lines indicate the start of each DSS cycle. (B) Body weight evolution during the acute DSS protocol, expressed as percentage of day 0. (C) Disease activity index (DAI) during acute colitis. (D) Colon length measured at sacrifice after the acute DSS cycle. (E) Blinded histopathological scores after acute colitis. (F) Body weight evolution throughout the two-cycle DSS protocol, expressed as percentage of day 0. (G) DAI during chronic colitis. (H) Colon length measured at sacrifice after the chronic DSS cycle. (I) Blinded histopathological scores after chronic colitis. (J) Ratio of *Tbx21* and *Foxp3* transcript levels in colonic tissue after the two-cycle DSS protocol, expressed as fold change relative to CD4-WT. (K) RT–qPCR analysis of epithelial barrier–associated transcripts in colonic tissue after the two-cycle DSS protocol, expressed as fold change relative to CD4-WT. (L) Schematic of the azoxymethane (AOM)/DSS colitis-associated colorectal cancer model in CD4-WT, CD4-KO, and CD4-OE mice (AOM 10 mg/kg, i.p., followed by three cycles of 2% DSS for 5 days each, separated by recovery periods; colons collected at day 70). (M) Number of macroscopic tumors per colon. (N) Tumor burden expressed as percentage of colon area occupied by tumors. (O) Flow cytometric analysis of tumor-infiltrating CD4^+^ T cells, showing IFNγ and IL-10 expression profiles. Data are shown as mean ± SEM (n = 3–11 mice per genotype in the acute DSS model; n = 7–15 mice per genotype in the chronic DSS model; n = 7–12 mice per genotype for tumor analysis pooled from two independent experiments; and n = 4–8 mice per genotype for flow cytometry). Statistical significance was assessed using two-way ANOVA with Tukey’s post hoc test for longitudinal measurements and one-way ANOVA with Tukey’s multiple-comparison test for endpoint analyses; *p < 0.05, **p < 0.01, ***p < 0.001; trends (p < 0.1) are indicated where applicable. Endpoint: e.p. Intraeritoneal: i.p.

### 4. Increasing T cell–intrinsic lamin A/C levels shifts the effector–regulatory balance and aggravates acute and chronic colitis

To assess the role of lamin A/C within T cells during colitis, mouse models with T cell–specific lamin A/C knockout (CD4-KO) or lamin A overexpression (CD4-OE) were challenged with one- or two-cycle DSS protocols to model acute and chronic disease, respectively (Fig. 3A). In the acute one-cycle DSS model, CD4-KO mice exhibited reduced body weight loss and lower DAI, without significant differences in colon length and histological damage score compared with CD4-WT controls or CD4-OE (Fig. 3B–E). In contrast, CD4-OE mice did not show significant differences relative to CD4-WT controls in body weight loss, DAI, colon length, or histological damage score, indicating no consistent effect on acute colitis severity (Fig. 3B–E). In the two-cycle DSS model, disease severity followed a genotype-dependent gradient, with CD4-KO mice showing the mildest phenotype, CD4-WT mice an intermediate phenotype, and CD4-OE mice the most severe phenotype (Fig. 3F–I). Compared with CD4-WT controls, CD4-KO mice exhibited reduced body weight loss and lower DAI, whereas CD4-OE mice showed greater body weight loss, higher DAI, reduced colon length, and increased histological damage. In addition, CD4-OE mice displayed greater body weight loss, higher DAI, and more pronounced colon shortening than CD4-KO mice (Fig. 3F–I). In this two-cycle DSS model, transcriptional analysis of colonic tissue revealed an increased *Tbx21* (T-bet)/*Foxp3* ratio in CD4-OE mice compared with CD4-KO mice, indicating a shift toward effector-dominated responses (Fig. 3J). No consistent genotype-dependent differences were detected for transcripts associated with epithelial barrier integrity (Fig. 3K). These findings indicate that T cell–intrinsic lamin A/C levels reprogram the effector–regulatory balance, with reduced lamin A/C favoring protection and overexpression promoting more severe chronic colitis.

### 5. Reducing T cell lamin A/C levels protects against colitis-associated colorectal cancer by shifting the effector–regulatory balance

IBD is associated with an increased risk of colorectal cancer (CRC), driven both by chronic, sustained inflammation [30] and by changes in T cell–mediated immune surveillance [31] among other contributory mechanisms. As T cell–specific lamin A/C modulation altered the effector/regulatory balance and intestinal inflammation in our models, we investigated whether these changes translated into altered susceptibility to colitis-associated CRC. In a colitis-associated CRC model induced by AOM followed by repeated DSS cycles (Fig. 3L), disease evolution mirrored colitis severity with exacerbated pathology in CD4-OE mice and partial protection in CD4-KO mice (Suppl. Fig. 4 A-C). Tumor burden was strongly genotype dependent: CD4-KO mice developed very few tumors, whereas CD4-OE mice showed increased tumor number and area, with WT animals displaying intermediate values (Fig. 3M, N; Suppl. Fig. 4 D). Immunophenotyping revealed that tumors from CD4-OE mice were enriched in IFNγ-producing CD4⁺ T cells, whereas tumors from CD4-KO mice showed higher frequencies of IL-10–producing CD4⁺ T cells populations, consistent with a more regulatory tumor microenvironment (Fig. 3O). These findings indicate that T cell lamin A/C levels influence tumor development by shaping the effector–regulatory T cell balance.

### 6. Fibroblast-specific lamin A/C deletion establishes a tissue-protective niche and attenuates colitis

The contribution of fibroblast-derived stromal cells to intestinal inflammation was investigated using fibroblast-specific lamin A/C knockout mice (Col1a2-KO) and WT controls (Col1a2-WT). Mice were first treated with tamoxifen to induce lamin A/C deletion and then subjected to one- or two-cycle DSS protocols (Fig. 4A). In the one-cycle DSS model, Col1a2-KO mice showed reduced body weight loss compared with Col1a2-WT controls, whereas DAI and colon length were not consistently different (Fig. 4B–D). Histological damage was markedly reduced in Col1a2-KO mice compared with Col1a2-WT controls (Fig. 4E). In the two-cycle DSS model, protective effects were more pronounced, as Col1a2-KO mice exhibited lower DAI, reduced colon shortening, and significantly attenuated histological injury compared with Col1a2-WT controls (Fig. 4B, C, F, G). Transcriptional profiling demonstrated that fibroblast-specific lamin A/C deletion induced a tissue-protective program. After one DSS cycle, Col1a2-KO mice showed increased expression of immunoregulatory cytokines (*Il10*, *Il22*, *Tgfb1*) and epithelial barrier-associated genes (*Muc1*, *Marveld2*, *Tjp1*) compared with Col1a2-WT controls. Elevated *H2-Ab1 (MHC-II)* and *Stat3* expression, together with a trend toward higher *Tbx21* levels, indicated early mucosal remodeling (Fig. 4H). Following two DSS cycles, a core protective signature persisted, with sustained upregulation of *Il10*, *Muc1*, and *Marveld2*, reflecting stabilization of regulatory and epithelial-supportive programs (Fig. 4I). Flow cytometry of mesenteric lymph nodes revealed reduced frequencies of activated effector-like CD4⁺ Th17 cells (CD4^+^RORγt^+^) and a tendency toward increased Treg frequencies (CD4^+^CD25^+^Foxp3^+^) (Fig. 4J), indicating that stromal lamin A/C deletion limits peripheral T cell effector differentiation. Collectively, these results identify fibroblast lamin A/C as a key regulator of the stromal-immune interface, whose loss reinforces epithelial integrity and establishes a tissue-protective niche.

**Figure 4.**
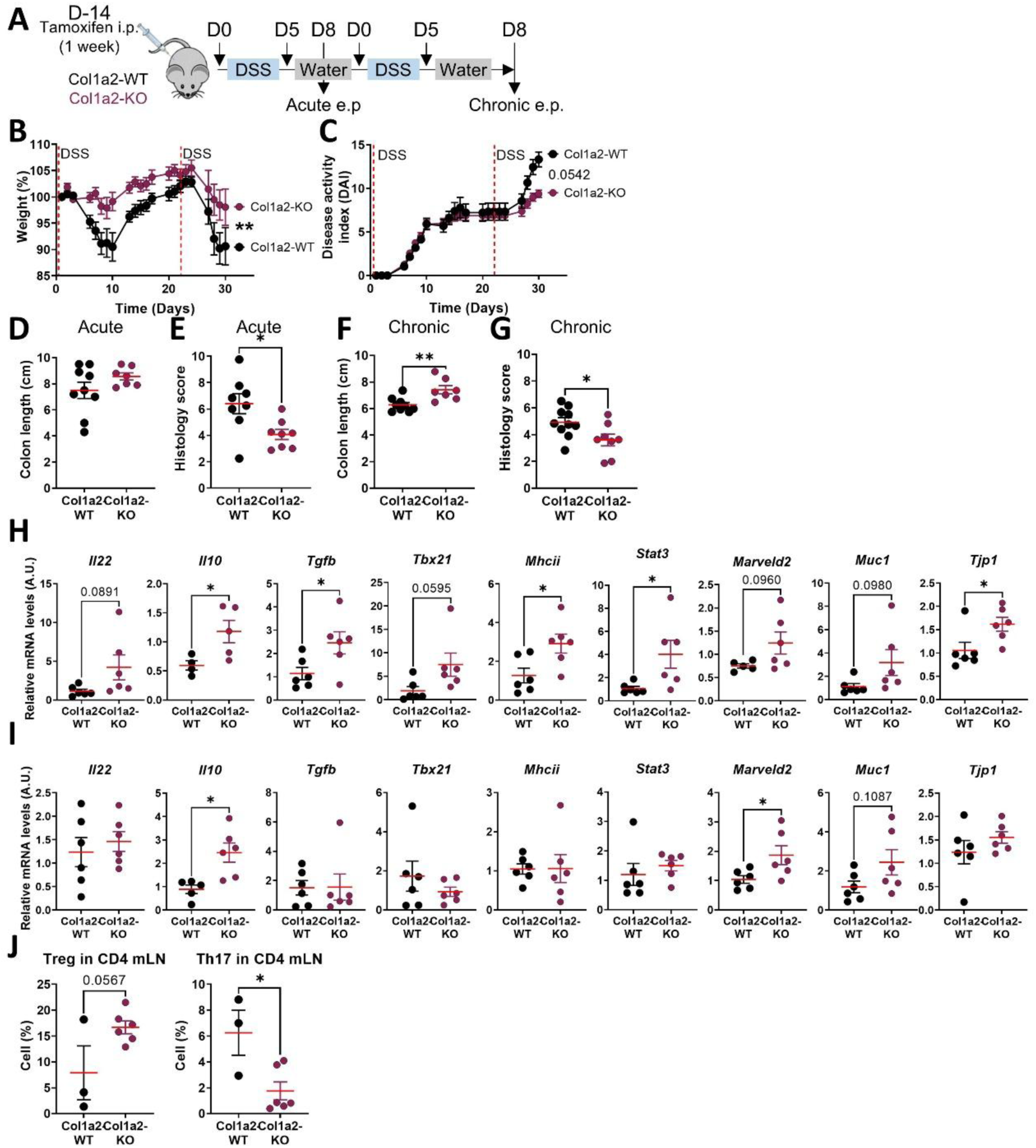
Fibroblast-specific lamin A/C modulates acute and chronic DSS-induced colitis by promoting a tissue-protective transcriptional and cellular program. (A) Schematic representation of the DSS-induced colitis protocols used to assess the role of fibroblast-intrinsic lamin A/C in wild-type (Col1a2-WT) and fibroblast-specific lamin A/C-deficient (Col1a2-KO) mice, using one-cycle acute DSS colitis or two-cycle chronic DSS colitis. Discontinuous vertical lines indicate the start of each DSS cycle. Mice were treated with tamoxifen (i.p.) for 1 week, 2 weeks before DSS initiation, to induce Cre recombination. (B) Body weight evolution during the acute and chronic DSS protocols, expressed as percentage of day 0. (C) Disease activity index (DAI) during DSS treatment. (D) Colon length measured at sacrifice after the acute DSS cycle. (E) Blinded histopathological scores after the acute DSS cycle. (F) Colon length measured at sacrifice after the chronic DSS cycle. (G) Blinded histopathological scores after the chronic DSS cycle. (H) RT-qPCR analysis of the indicated transcripts in colonic tissue after the acute DSS cycle, expressed as fold change relative to Col1a2-WT. (I) RT-qPCR analysis of the indicated transcripts in colonic tissue after the chronic DSS cycle, expressed as fold change relative to Col1a2-WT. (L) Flow cytometric analysis of mesenteric lymph nodes (mLN) after the chronic DSS model, assessing Treg (CD4^+^CD25^+^Foxp3^+^) and Th17 (CD4^+^RORγt^+^) T-cell populations among CD4^+^ cells. Data are shown as mean ± SEM (n = 6–9 mice per genotype). Statistical significance was assessed using two-way ANOVA with Tukey’s post hoc test for longitudinal measurements and unpaired Student’s t-test for endpoint analyses; *p < 0.05, **p < 0.01; trends (p < 0.1) are indicated where applicable. Endpoint: e.p. Intraperitoneal: i.p.

### 7. T-cell Lamin A/C levels associate with severe histological damage in human IBD

To assess whether the compartment-specific lamin A/C programs identified in mice are reflected in human disease, lamin A/C levels were quantified by immunostaining in CD3⁺ T cells from formalin-fixed, paraffin-embedded (FFPE) intestinal biopsies of patients with IBD (N=16), including CD (N=8) and UC (N=8) and non-IBD controls (n=10) (Supp. Fig. 5, Fig. 5A-C, Supp. Table 2). Lamin A/C expression was modeled against CD3 abundance, perinuclear area as a proxy for nuclear expansion, and disease status using linear mixed-effects models (Supp. Table 3-5). In non-IBD controls, lamin A/C expression increased in parallel with higher CD3 abundance and larger perinuclear area in lamina propria CD3^+^ T cells (p < 0.001) (Supp. Table 4, Fig. 5B). However, this upregulation attenuated in the IBD microenvironment, with significant negative interactions between IBD status and both CD3 abundance and perinuclear area (p < 0.001) (Supp. Table 4, Fig. 5B). Stepwise selection identified overweight as a marginal covariate trending toward reduced lamin A/C (p = 0.051, Supp. Table 3-4, Supp. Figure 6A). This attenuated lamin A/C pattern was conserved across both disease subtypes (Supp. Table 4, Supp. Fig. 6B). Furthermore, substituting IBD status with a continuous histological severity score yielded an identical pattern: local tissue damage severity negatively interacted with both CD3 abundance and nuclear expansion (p < 0.001) (Supp. Table 4, Fig. 5C). Collectively, these data corroborate our murine models, demonstrating that intestinal inflammation disrupts the dynamic regulation of lamin A/C in lamina propria T cells. These findings suggest that T-cell lamin A/C levels reflect disease severity rather than disease presence.

**Figure 5.**
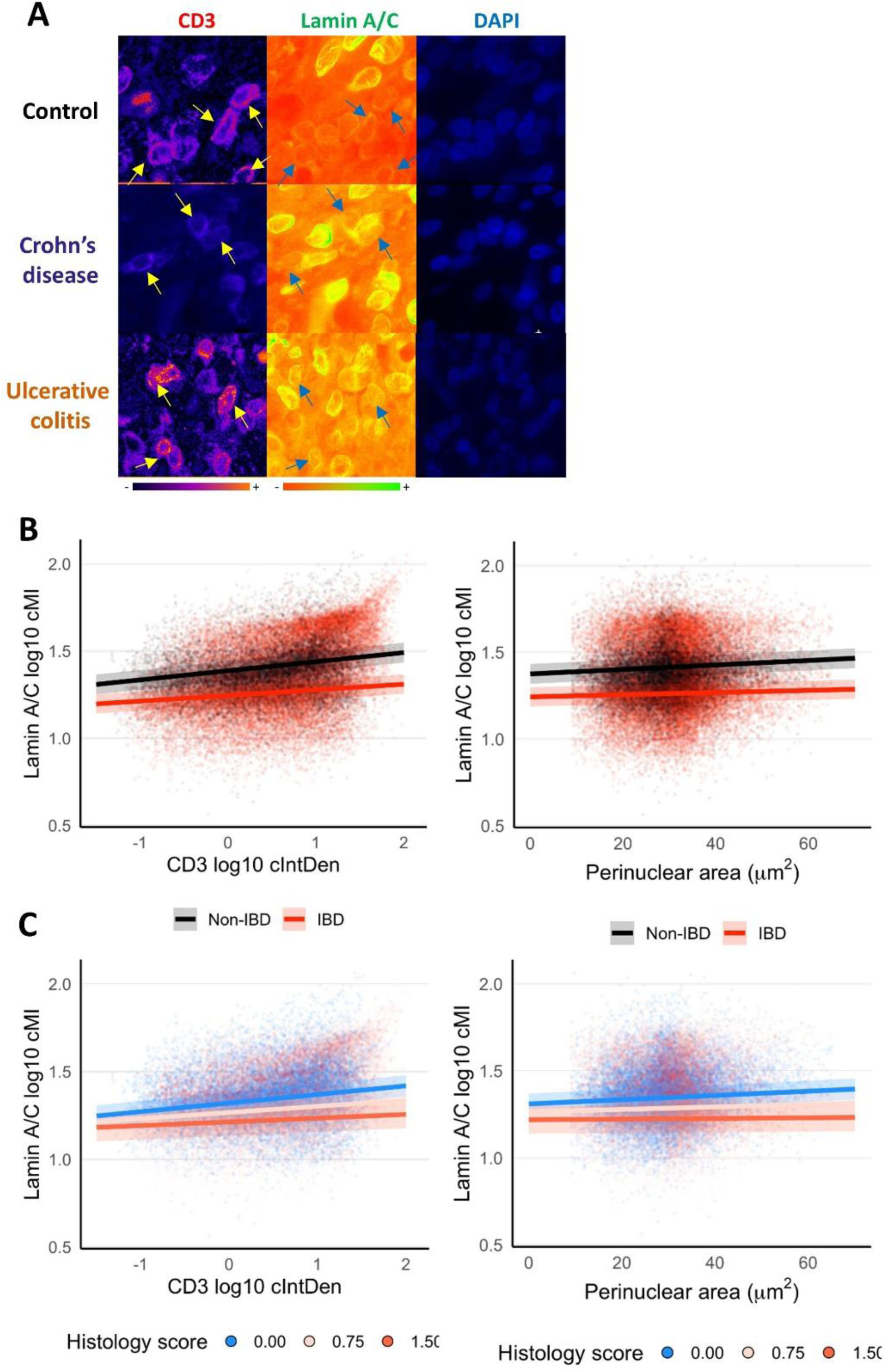
Lamin A/C Expression in CD3^+^ Cells in Human IBD. A) Representative immunofluorescence images in pseudocolor of lamina propria areas from FFPE intestinal samples from non-IBD controls and patients with Crohn’s disease (CD) or ulcerative colitis (UC) co-stained for CD3, lamin A/C, and DAPI. Yellow and blue arrows highlight representative CD3^+^ cells. Color scales indicates CD3 and lamin A/C signal intensity. B) Lamin A/C corrected mean intensity (cMI) vs CD3 corrected integrated density (cIntDen) or perinuclear area in the primary mixed-effects model according to IBD status. C) Lamin A/C cMI vs CD3 cIntDen or perinuclear area in the secondary mixed-effects model according to histological severity score. Colored lines represent model fits, and shaded areas indicate confidence intervals.

**Figure 6.**
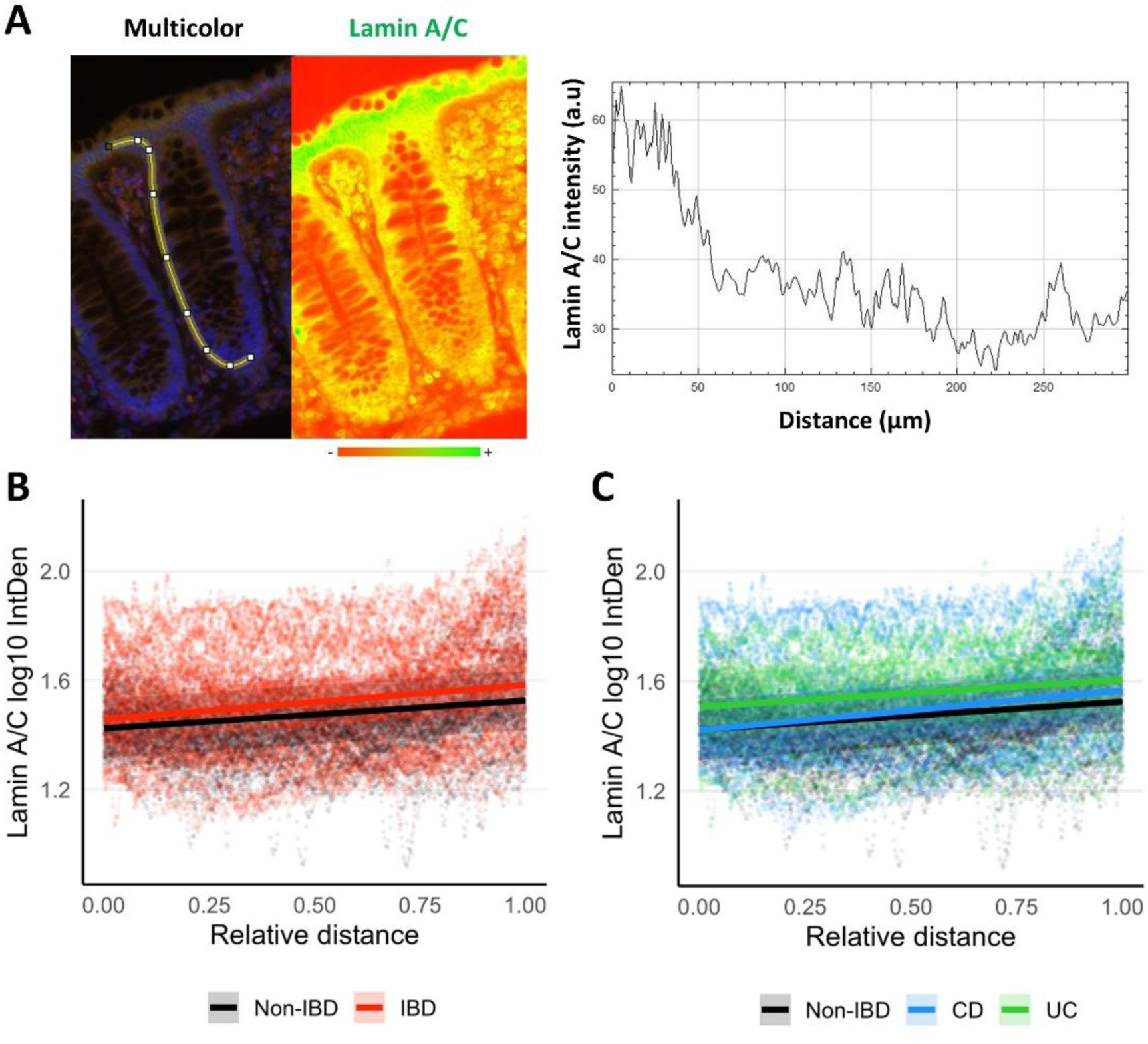
Lamin A/C Expression in Intestinal Crypts in Human IBD. (A) Representative immunofluorescence image of an intestinal crypt stained for lamin A/C and DAPI. The left panel shows the multicolor image overlaid on the epithelial crypt region of interest; the middle panel shows lamin A/C in pseudocolor for improved visualization of signal differences; and the right panel shows the fluorescence intensity profile along the marked line, with distance 0 corresponding to the top of the crypt and increasing distance extending toward the basal side. Color scale indicates lamin A/C signal intensity. B) Lamin A/C integrated density (IntDen) vs relative distance from the crypt base to the luminal surface in the primary mixed-effects model according to IBD status. C) Lamin A/C IntDen versus relative distance from the crypt base to the luminal surface in the secondary mixed-effects model according to IBD subtype.

### 8. Epithelial lamin A/C spatial gradient is altered in human IBD

Lamin A/C distribution within the intestinal epithelium was quantified along the crypt axis in FFPE samples from non-IBD, UC, and CD patients (Fig. 6A) and analyzed using mixed-effects models (Supp. Table 6-8). To account for significant IBD-associated crypt hyperplasia (Supp. Fig. 7A), crypt depth was normalized to a relative base-to-lumen distance. In non-IBD controls, lamin A/C exhibited a robust spatial increase toward the lumen (p < 0.001; Fig. 6B-C, Supp. Table 7), and patient age was identified as an independent covariate associated with lower overall expression (p = 0.032; Supp. Fig. 7B-D, Supp. Table 6-7). Modeling also revealed a significantly steeper spatial gradient of lamin A/C in IBD than in controls (interaction p < 0.001; Fig. 6B, Supp. Table 7). Subtype stratification showed that this accelerated trajectory was driven entirely in CD (p < 0.001); whereas the gradient in UC was indistinguishable from controls (p = 0.278) (Fig. 6C, Supp. Table 7). Furthermore, the epithelial gradient did not correlate with histological severity (p = 0.389; Supp. Fig. 7D, Supp. Table 7). These data suggest that altered epithelial lamin A/C spatial regulation is not a generalized inflammatory response, but rather a pathology-specific feature of CD.

## Discussion

Our study identifies lamin A/C as a compartment- and context-dependent regulator of intestinal inflammation and colitis-associated tumorigenesis. Across myeloid cells, T cells, fibroblasts, and the intestinal epithelium, lamin A/C exerts distinct functions that converge on shared disease outcomes in experimental colitis and colitis-associated CRC. Through genetic manipulation of lamin A/C in individual cellular lineages, we demonstrate that reduced lamin A/C expression in hematopoietic and stromal cells mitigates DSS-induced colitis, whereas increasing lamin A/C in T cells aggravates chronic inflammation and colitis-associated CRC. Within this framework, the innate immune compartment emerges as a major site of lamin A/C-dependent regulation. Within the innate immune compartment, our data identify lamin A/C as a regulator of myeloid pathogenicity in colitis. Myeloid-specific lamin A/C deletion attenuated DSS colitis, with a clearer divergence in the chronic than in the acute setting, supporting the idea that lamin A/C-dependent myeloid programs contribute not only to early innate inflammation but also to the sustained immune circuits that maintain tissue damage. In line with this, adoptive transfer of lamin A/C-deficient BMDCs was sufficient to recapitulate protection and was associated with increased Treg frequencies and reduced IFNγ-producing CD8⁺ T cells, consistent with altered antigen-presenting cell function. Our proteomic profiling of LysM-KO BMDCs provides a mechanistic framework for this phenotype, revealing suppression of pathways linked to antigen uptake, processing, and presentation together with enrichment of lipid/carbohydrate metabolism and mucosal defense programs. These findings are consistent with a shift toward a less inflammatory and less immunogenic myeloid state in vivo. This interpretation fits with prior studies showing that macrophage lamin A/C can promote NF-κB-driven inflammatory programs and contributes to obesity-associated metabolic inflammation [32]. More broadly, lamin A/C has also been linked to NF-κB regulation through chromatin-associated mechanisms, including Ser22-phosphorylated lamin A/C localization to the nuclear interior, association with active enhancers, and effects on NF-κB nuclear localization and gene accessibility [33, 34]. It is also consistent with the role of lamin A/C in DC functions required for efficient priming of Th1 and cytotoxic responses, including synapse-related processes and inflammatory signaling [20]. Rather than assigning a single downstream pathway, our data support a model in which lamin A/C sustains inflammatory and antigen-presenting myeloid states that can amplify both direct innate effector functions, such as cytokine and chemokine production and pattern-recognition receptor-driven inflammatory signaling affecting epithelial barrier integrity [35], and indirect effects on adaptive immunity through antigen-presenting cell -dependent T-cell shaping in the gut [36]. At the same time, lamin A/C biology appears to be highly context-dependent across myeloid settings. In macrophages, pro-inflammatory activation can trigger lamin A/C phosphorylation and degradation, augmenting IFN-β expression and downstream STAT signaling, whereas pharmacological blockade of lamin A/C loss dampens inflammatory gene expression [21]. In addition, physical context may further tune these responses, as mechanical microenvironments can modulate lamin A/C abundance and link the nuclear lamina to actin-dependent transcriptional programs that shape macrophage inflammatory outputs [37, 38]. A broader relevance across myeloid lineages is also supported by evidence that lamin A/C deficiency suppresses LPS-driven activation phenotypes in microglia and highlights STAT1-centered networks as candidate downstream hubs in inflammatory activation programs [39]. Taken together, these considerations argue against extrapolating a single universal lamin A/C-dependent mechanism across all inflammatory settings, while still supporting the conclusion that, in DSS colitis, the net effect of constitutive myeloid lamin A/C loss is protective and is likely mediated in substantial part by dampened inflammatory and antigen-presenting functions.

In the adaptive immune compartment, our results identify lamin A/C as an important regulator of effector–regulatory balance and a key determinant of chronic T cell–driven pathology in our models. T cell-specific lamin A/C overexpression exacerbated chronic DSS colitis and shifted colonic transcriptional readouts toward an effector-dominant state, as reflected by the increased *Tbx21*/*Foxp3* ratio, consistent with prior mechanistic work showing that lamin A/C promotes Th1 differentiation and restrains Treg programs [26, 27]. Extending these findings to tumorigenesis, T cell-specific lamin A/C deletion markedly reduced tumor burden in the AOM/DSS model, in line with the established link between sustained colitis severity and colitis-associated cancer risk [40] and with the broader concept that the immune microenvironment shapes CRC evolution [31]. By contrast, although lamin A/C overexpression aggravated chronic inflammation, this did not translate into a further increase in tumor burden relative to WT, suggesting that the relationship between T cell inflammatory activity and tumor promotion is not strictly linear. One possible explanation is that IFNγ-associated effector programs, while contributing to tissue damage, may also preserve elements of anti-tumor immune surveillance in some contexts [41].

Consistent with this interpretation, tumors from CD4-OE mice were enriched in IFNγ-producing CD4 T cells, whereas CD4-KO tumors showed a more regulatory IL-10-associated profile. Together, these results place lamin A/C at the intersection of T cell differentiation, chronic tissue injury, and inflammation-associated tumor development.

The stromal compartment emerged as an important site of lamin A/C-dependent regulation in our models. Fibroblast-specific lamin A/C deletion conferred significant protection in DSS colitis and was associated with increased expression of Il10, Tgfb1, Il22 and epithelial barrier-associated genes, consistent with a tissue-protective stromal program and with the established capacity of intestinal stromal cells to regulate mucosal immunity and epithelial homeostasis [11, 42]. Our immune profiling further supports the view that a lamin A/C-deficient stromal niche can favor a more regulatory local environment, as reflected by reduced Th17-like signatures and a trend toward increased Tregs in lymphoid compartments, in agreement with prior evidence that colonic myofibroblasts can promote Treg expansion [43]. Although the fibroblast-intrinsic molecular pathways downstream of lamin A/C remain to be defined, these findings identify lamin A/C as a determinant of stromal functional state, with cell-extrinsic consequences for epithelial barrier support and adaptive immune regulation.

The translational relevance of these compartment-specific mechanisms is supported by our analyses of human IBD biopsies, which revealed lamin A/C patterns broadly consistent with the murine findings in a compartment-resolved manner. In CD3⁺ T cells from intestinal biopsies, lamin A/C expression increased with T-cell abundance and perinuclear area, as a proxy for nuclear expansion, in non-IBD controls, but this adaptive upregulation was blunted in IBD, with significant negative interactions for both CD3 and nuclear expansion. Overweight status also tended to associate with lower lamin A/C, and the same interaction pattern was recapitulated when IBD diagnosis was replaced by a continuous histological severity score, indicating that local tissue damage modulates the lamin A/C response in infiltrating T cells. These observations, together with our genetic models in mice, support a view in which T-cell lamin A/C levels report the intensity and quality of the inflammatory microenvironment—integrating immune activation, nuclear remodeling, and patient-related factors such as overweight—rather than merely indicating the presence or absence of IBD.

Beyond the immune infiltrate, we quantified lamin A/C specifically in epithelial cells along the crypt axis, revealing a spatial gradient that increased from the crypt base to the luminal surface. In non-IBD controls, epithelial lamin A/C increased toward the lumen, consistent with prior studies showing higher lamin A/C levels in differentiated epithelial cells than in stem/progenitor cells along the crypt axis [44, 45]. Patient age emerged as an independent covariate associated with lower overall epithelial lamin A/C levels. This age-related decline resonates with reports that lamin A/C expression decreases in hematopoietic and immune cells during human aging, with functional consequences for vascular and immune homeostasis [46]. In IBD, the epithelial gradient was significantly steeper than in controls, indicating a sharper increase in lamin A/C toward the luminal/top crypt compartment. This change was driven by CD, whereas UC largely preserved the control-like pattern. This CD-specific steepening occurs in the context of pathological epithelial remodeling, including de novo keratin 7 expression linked to drug resistance and cytoskeletal reorganization [47], and aligns with the concept that cytoplasmic keratins couple with and maintain nuclear envelope integrity in colonic epithelial cells [48]. Together, these data suggest that altered epithelial lamin A/C spatial regulation is not a generalized inflammatory response, but rather a pathology-specific feature of CD that reflects fundamental rewiring of the crypt differentiation program.

Taken together, our murine and human data identify lamin A/C as a compartment-and context-dependent regulator whose net impact on IBD reflects the integrated output of immune, stromal, and epithelial programs. In this framework, T-cell lamin A/C levels emerge as a readout of mucosal inflammatory burden, while epithelial lamin A/C gradients capture CD-specific epithelial stress, providing complementary information to conventional histology within the same biopsy.

Collectively, these results support a model in which lamin A/C does not operate through a single unified program shared across compartments but instead shapes lineage-specific pathogenic or protective programs whose combined output determines the net balance between tissue injury, repair, and tumor-promoting inflammation.

These findings raise the possibility that quantitative assessment and targeted modulation of lamin A/C across intestinal compartments could inform patient stratification and guide therapeutic strategies aimed at dampening pathogenic inflammation while preserving epithelial protection and limiting the risk of colitis-associated CRC.

## Materials and Methods

### Ethics approval

Human studies complied with the Declaration of Helsinki and were approved by the Auria Biobank (Turku, Finland) [47]. Animal experiments followed EU Directive 2010/63/EU and Spanish law, approved by the CNIC Animal Care Committee and the Spanish Ministry (PROEX 203/17, 100.4/22, 244.1/22).

### Human tissue samples

FFPE intestinal resection specimens from CD, UC, and non-inflammatory controls were obtained from the Auria Biobank. A pathologist selected areas with active lamina propria inflammation. Clinical and histological data were collected [47].

### Mice

Mice were bred and maintained under specific pathogen-free (SPF) conditions at the CNIC and Imas12. All strains were on a C57BL/6 background. Wild-type (C57BL/6-CD45.1 or CD45.2) mice were obtained from the CNIC internal colony or from Jackson Laboratory. The following conditional genetically modified strains were generated and used: Vav1-Cre [49] × Lmna^flox/flox^ [50] (Vav1-KO) for pan-hematopoietic deletion of lamin A/C, and Vav1-Cre × Lmna^OE^ [46] (Vav1-OE) for pan-hematopoietic overexpression of lamin A; Vav1-Cre × Lmna^flox/flox^ × Rag1^-/-^ [51] (Vav1-KO-Rag1) for pan-hematopoietic deletion of lamin A/C in the absence of T and B cells; LysM-Cre [52] × Lmna^flox/flox^ (LysM-KO) for myeloid-specific deletion of lamin A/C; CD4-Cre × Lmna^flox/flox^ (CD4-KO) for T cell–specific deletion of lamin A/C, and CD4-Cre [53] × Lmna^OE^ (CD4-OE) for T cell–specific overexpression of lamin A; and Col1a2-CreERT2 [54] × Lmna^flox/flox^ (Col1a2-KO) for tamoxifen-inducible, fibroblast-specific deletion of lamin A/C. Rag1^-/-^ [51] mice and Cre-negative littermates served as controls. Female mice aged 8–12 weeks were used, and experiments were performed with sex-matched animals. For fibroblast-specific deletion, tamoxifen (Sigma-Aldrich) was administered intraperitoneally (i.p.) at 1 mg per mouse daily for 5 consecutive days, two weeks prior to colitis induction.

#### Induction of colitis and colitis-associated cancer

For acute DSS colitis, mice received 2% DSS (MW ∼40,000; Alfa Aesar) in drinking water for 5-6 days and were euthanized on day 8-10, except for Vav1-KO-Rag1 mices, which received 1% DSS under the same conditions. Chronic colitis involved a second 2% DSS cycle after recovery. For colitis-associated CRC, mice received intraperitoneal azoxymethane (AOM; 10 mg/kg; Merck), followed one week later by three 5-day cycles of 2% DSS, and were euthanized on day 70 for tumor assessment. Disease activity was scored daily as previously described [55]: stool consistency: 0 (normal), 1 (loose stool), and 2 (diarrhea); rectal prolapse: 0 (absent) and 1 (present); rectal bleeding: 0 (absent), 1 (low), and 2 (pronounced); and spine curvature: 0 (absent), 1 (low), and 2 (pronounced); mouse activity: 0 (normal), 1 (low), 2 (absence of movement).

#### Histopathological analysis

Colons were fixed, embedded, and 5 µm sections were hematoxylin and eosin stained. Inflammation was scored (0–3) based on leukocyte infiltration, epithelial hyperplasia, submucosal inflammation, crypt damage, goblet cell depletion, crypt abscesses, and lymphocyte accumulation. An average score from 2-4 blinded sections per mouse was calculated [56].

#### Immune cell isolation and flow cytometry

Colon immune cells were isolated using an adapted protocol [57]. Cleaned colons were opened longitudinally, cut into ∼0.5 cm fragments, and incubated in intraepithelial lymphocyte (IEL) medium consisting of phosphate-buffered saline (PBS) supplemented with 10% FBS (Sigma-Aldrich), 5 mM EDTA (Sigma), and 15 mM HEPES (Hyclone) (30 ml/colon) for 30 min at 37 °C with agitation (200 rpm). Intraepithelial cells were recovered from the supernatant after vigorous shaking, filtration (40 µm), and centrifugation (350 g, 10 min). Remaining tissue was washed to remove EDTA and digested in lamina propria cell (LPC) medium consisting of RPMI supplemented with 10% FBS (Sigma-Aldrich) and 15 mM HEPES (Hyclone), with collagenase VIII (Sigma) added immediately before use to a final concentration of 0.4 mg/ml (300 U/ml), for 45 min at 37 °C with agitation. Lamina propria cells were obtained after filtration and centrifugation. Cells were stained with fluorophore-conjugated antibodies (Supplementary Table 1), fixed/permeabilized when required wit BD cytofix/cytoperm and prem/wash solucitons for cytokine staining, and Invitrogen FoxP3/Transcription Factor staining buffer set for nuclear stainings, acquired on BD or Cytek cytometers, and analyzed using FlowJo v10.

#### Bone marrow-derived dendritic cell (BMDC) generation and adoptive transfer

BMDCs were generated from bone marrow precursors cultured with GM-CSF for 7–9 days and matured by incubation with the TLR4 agonist lipopolysaccharide (LPS) from Escherichia coli O111:B4 (20 ng/mL; Sigma-Aldrich). Following a standard DSS induction protocol, WT mice were randomly assigned to two groups receiving BMDCs derived from WT or LysM-KO donors. BMDC transfer experiments were performed following an established protocol adapted from [58], with minor modifications. Three days prior to DSS exposure, mice were injected i.p. with 2 × 10^6^ WT or LysM-KO BMDCs in a minimal volume of sterile, endotoxin-free PBS. Animals were then provided *ad libitum* access to drinking water containing 2% DSS for 5 days. On day 3 of DSS exposure, a second intraperitoneal injection of 2 × 10^6^ BMDCs was administered. Mice were euthanized 10 days after the initiation of DSS treatment.

#### RNA extraction and quantitative PCR

Comparable tissue sections (<30 mg) from the same intestinal region were collected from each mouse and homogenized using a TissueRuptor II (Qiagen) in either TRI Reagent (Invitrogen) or lysis buffer from the RNeasy Mini Kit (Qiagen). Total RNA was isolated according to the manufacturer’s instructions, and RNA concentration and purity were assessed by measuring absorbance at 260/280 nm using a NanoDrop One spectrophotometer (Thermo Scientific). Total RNA (0.5–2 μg) was reverse transcribed using the High-Capacity cDNA Reverse Transcription Kit with RNase Inhibitor (Applied Biosystems). The resulting cDNA was diluted 1:100, and quantitative PCR was performed in technical duplicates using Power SYBR Green PCR Master Mix (Applied Biosystems) on a 7500 Fast Real-Time PCR System (Applied Biosystems). Gene expression was normalized to stably expressed housekeeping genes (*Actb*, *Gapdh*, *Hprt1*, *Ywhaz*, or *18s*), and relative expression levels were calculated using the 2−ΔΔCt method, with the wild-type group used as the reference control. Primer sequences are listed in Table M3.

#### Human tissue samples and histopathological evaluation

FFPE intestinal specimens from IBD patients and non-IBD controls were obtained from the Auria Biobank (Turku, Finland) [47], and processed in collaboration with the laboratory of Diana Toivola at Åbo Akademi University, where samples were stained and analyzed, and with Prof. Markku Voutilainen at University of Turku Hospital. IBD samples were collected during colectomy or ileal resection from colonic regions between the sigmoid and ascending colon in patients with either CD or UC. Non-IBD control biopsies were harvested during ileocolonoscopy from individuals with suspected but subsequently discarded IBD or colorectal cancer diagnoses. These patients presented with symptoms such as prolonged diarrhea, anemia, gastrointestinal bleeding, or weight loss. Transport, handling, and storage of the paraffin-embedded tissue samples were carried out according to biobank standard operating procedures. Patient demographics, clinical characteristics and treatment were also collected, and all data were anonymized. Histological severity was independently assessed by a pathologist across two selected regions of interest (ROI) per patient. Six hallmark inflammatory features (changes in epithelium, lamina propria neutrophils, epithelial neutrophils, erosions/ulcers, general inflammatory activity, and granulomas) were visually graded on a standard ordinal scale. To facilitate quantitative analysis, a composite mean histology severity score (ranging from 0 to 3) was calculated for each region of interest (ROI) to represent the aggregate tissue damage [47].

#### Immunofluorescence of human samples

FFPE sections were incubated at 65 °C for 30 min, deparaffinized in xylol, rehydrated through graded ethanol, and subjected to antigen retrieval in 10 mM citrate buffer (pH 6.0) for 10 min at 95–100 °C. Autofluorescence was quenched by incubation with 100 mM glycine for 1 h at room temperature. Sections were blocked with PBS containing 5% normal goat and donkey serum and stained overnight at 4 °C with rabbit anti-human CD3 (1:100; DAKO). After washing and permeabilization (PBS with 0.3% Triton X-100), intracellular staining was performed using anti–lamin A/C–Alexa Fluor 488 (1:50; Cell Signaling) overnight at 4 °C. Slides were then incubated with donkey anti-rabbit Alexa Fluor 546 (1:200; Invitrogen) for 1 h, counterstained with DAPI and DRAQ5, and mounted with ProLong Gold (Thermo Fisher Scientific). Sample order was randomized to mitigate potential batch effects between groups, given that not all could be processed at once. Slides were scanned using a Pannoramic Midi fluorescence slide scanner (3DHISTECH Ltd.) at 40× magnification.

#### Immunofluorescence analysis of lamin A/C in CD3⁺ cells and crypt epithelium in human samples

Digital whole-slide images were imported into QuPath v0.4.4 [59]. From each of the two pathologist-selected ROIs per patient, sub-regions enriched in CD3⁺ cells within the lamina propria were selected and exported to Fiji/ImageJ [60] as multichannel composite images. Nuclear segmentation was performed on the nuclear channel using a StarDist-based workflow. Because the nuclear lamina lies directly apposed to the inner nuclear envelope and T cells possess minimal cytoplasm, we defined a standardized spatial geometry to capture both markers: a perinuclear ring mask extending 5 pixels inward and 5 pixels outward from the segmented nuclear membrane boundary. CD3 spatial segmentation utilized a minimum intensity threshold to prevent signal saturation and standardize foreground detection across optical variations, whereas no threshold was used for the lamin A/C signal to capture the continuous structural envelope. Background fluorescence was measured in tissue-free regions and subtracted from all measurements. To rigorously identify true T cells and exclude artifacts, stringent geometric and signal filters were applied. Cells were excluded if they met any of the following criteria: lack of CD3 signal; a CD3 area < 5 µm² (spurious fluorescence); a perinuclear ring area > 65 µm² (potential incorrectly segmented cells); a CD3 area coverage (CD3 area divided by perinuclear ring area) < 5% (signal bleed-through from adjacent cells); or a CD3 area coverage > 99% (optical sectioning artifacts representing cells sliced near the poles). For lamin A/C quantification, we evaluated the corrected mean intensity (cMI; mean intensity minus local background) to assess changes in protein concentration within the perinuclear ring independent of cell size variations during activation. To proxy T-cell activation, CD3 total abundance was quantified using a threshold-adjusted corrected integrated density (cIntDen) to account for the protein’s discrete clustering. To prevent spatial bias and normalize for membrane expansion, cIntDen was calculated as the product of the threshold-adjusted mean intensity (MI minus minimum intensity threshold) and the CD3 area coverage. Both variables were log10-transformed for normalization. For intestinal crypt analysis, the same QuPath-selected ROIs were used. Epithelial regions encompassing complete crypts were manually delineated and exported to ImageJ. Using segmented line profiling, lamin A/C cIntDen was continuously measured along the crypt axis from the luminal surface to the crypt base. The relative distance from base was calculated to normalize crypt depth across all samples.

#### Quantitative proteomic analysis of BMDCs cells

BMDCs were generated from WT and LysM-KO mice by culturing bone marrow cells with GM-CSF for 7–9 days and matured overnight with LPS (100 ng/mL). Cells were harvested, lysed in 50 mM Tris-HCl with 2% SDS and 10 mM Tris(2-carboxyethyl) phosphine (TCEP), and protein concentration measured by Direct Detect IR spectrometry. Lysates were reduced, alkylated with iodoacetamide (IAA), digested with trypsin, and resulting peptides desalted on C18 cartridges. Peptides were labeled with TMT 10-plex reagents, fractionated by high-pH reversed-phase chromatography, and analyzed by LC-MS/MS on an Orbitrap Fusion mass spectrometer (Thermo Fisher Scientific). Protein identification was performed with SEQUEST-HT against the Uniprot mouse database with a 1% false discovery rate (FDR), and quantitative analysis was conducted with the SanXoT package based on the WSPP model [61, 62]. Proteins quantified by a single peptide, as well as known artifact proteins such as keratin, bovine serum, and trypsin contaminants were excluded for further analysis. Proteins with inconsistent expression between controls (i.e., top 1% with the highest within-group standard deviation) were removed to reduce noise and preserve statistical power. Within-group and between-group sample variability was assessed by principal component analysis (PCA). Statistical differences between groups were estimated using linear models from the R package “limma” [63] and volcanic plots. Gene set enrichment analysis (GSEA) was performed using the Kyoto Encyclopedia of Genes and Genomes (KEGG) database, and the R package “clusterProfiler” [64]. To enhance statistical power and ensure biological relevance, the KEGG database was curated pre GSEA. Drug development, organismal systems, human diseases, overview maps, non-mammalian biology, and other-tissue-specific categories were excluded. Exceptions were categories related to environmental adaptation, immune system/disease, bacterial infection, and colorectal cancer. Allowed set sizes were between 10 and 500, proteins were ranked by their limma-derived t-statistics, p values were adjusted by the Benjamini-Hochberg procedure, and a seed was set to ensure reproducibility. GSEA was visualized using CNET and ridgeline plots. The analyses were performed using R version 4.1.0 (2021-05-18).

#### Statistical analysis

Statistical analyses on murine data were performed using GraphPad Prism 9 (GraphPad Software, San Diego, CA, USA). Data normality was assessed with the Kolmogorov–Smirnov test. Normally distributed data are presented as mean ± standard error of the mean (SEM). Comparisons between two groups were performed using unpaired Student’s t-tests, while comparisons among multiple groups were conducted using one-way or two-way ANOVA with Tukey post hoc tests, as appropriate. Correlations between variables were assessed by simple linear regression. For human IF data, linear mixed-effects models were constructed using R version 4.1.0 (2021-05-18) with the lme4 package to account for hierarchical data structures and unbalanced repeated measures. For CD3^+^ cell analyses, lamin A/C log_10_ cMI was modeled with nested random effects (batch > patient > ROI > sub-region). Fixed effects included CD3 log_10_ cIntDen, perinuclear area, disease status (either IBD status, subtype, or histology score), patient-level characteristics (demographics, clinical characteristics, and treatment), and interaction terms to assess disease-specific lamin A/C scaling. The CD3 minimum intensity threshold was included as a technical covariate to statistically absorb slide-to-slide optical variations. For the crypt analysis, lamin A/C log_10_ cIntDen was modeled against the relative distance from the base, IBD status, and their interaction, adjusting for patient-level ROI covariates, with nested random effects (batch > patient > crypt). Final models were determined via stepwise forward selection, retaining variables based on minimizing the Akaike information criterion (AIC), assessing multicollinearity via the variance inflation factor (VIF), and confirming significance through likelihood ratio tests. To prevent overfitting from collinear patient-level variables, stepwise selection was restricted to the primary IBD models, while secondary models (subtype, histological score) retained only core predictors to ensure stability. Statistical significance was defined as p < 0.05, p < 0.01, p < 0.001, and p < 0.0001. Trends were considered for 0.05 ≤ p < 0.1 and represented with their actual p values; non-significant results were indicated as ns.

## Acknowledgments

R.G.-B. and M.O.-Z., as well as A.S. and J.-M.G.-G., contributed equally to this manuscript. We thank Yixian Zheng (Baltimore, MD) for providing the Lmna^fl/fl^ mouse; M.J. Andrés-Manzano (CNIC) for experimental assistance; Harri Kujari and Santeri Anttila (University of Turku) for their contributions to the patient IBD material; and Auria Biobank (Turku, Finland) for help in collecting clinical information and carrying out tissue sectioning.

## Author contributions

Conceptualization: JMGG, AS; Formal Analysis: JMGG, RGB, MOZ, BHF, AU, VF; Funding Acquisition: JMGG, AS, JLP, CS, FSM, DMT, GC, SMA, JV, VA; Investigation: RGB, MOZ, BHF, AU, LP, NM, SMB, GC, DMT, HSM, IMA, VZ, LP, JAL; Methodology MAP; Resources MAP, PG, VA, JLP, DMT, FSM, GC, SMA, LP; Visualization RGB, MOZ, BHF; Writing - Original Draft Preparation JMGG, AS, RGB, MOZ; Writing – Review & Editing JMGG, AS, RGB, MOZ, DMT, LP, FSM, VA, CS, GC.

## Data availability.

The mass spectrometry proteomics data have been deposited to the ProteomeXchange Consortium (http://proteomecentral.proteomexchange.org) via the PRIDE partner repository [65] with the dataset identifier PXD080116.

## Abbreviations

AOM –: Azoxymethane.
BMDC –: Bone marrow-derived dendritic cell.
CD –: Crohn’s disease.
CRC –: Colorectal cancer.
Cre –: Cre recombinase.
DAI –: Disease activity index.
DAPI –: 4′,6-diamidino-2-phenylindole.
DC –: Dendritic cell.
DSS –: Dextran sulfate sodium.
FDR –: False discovery rate.
FFPE –: Formalin-fixed, paraffin-embedded.
GM-CSF –: Granulocyte-macrophage colony-stimulating factor.
GO –: Gene Ontology.
IBD –: Inflammatory bowel disease.
IFN –: Interferon.
IFNγ –: Interferon gamma.
IL –: Interleukin.
i.p. –: Intraperitoneal/intraperitoneally.
KEGG –: Kyoto Encyclopedia of Genes and Genomes.
KO.: Knock out.
LC-MS/MS –: Liquid chromatography-tandem mass spectrometry.
LPS –: Lipopolysaccharide.
MHC-II –: Major histocompatibility complex class II.
MLN –: Mesenteric lymph node.
NF-κB –: Nuclear factor kappa B.
PCA –: Principal component analysis.
ROI –: Region of interest.
RT–qPCR –: Reverse transcription quantitative polymerase chain reaction.
SEM –: Standard error of the mean.
TGFβ –: Transforming growth factor beta.
Th1 –: T helper 1.
Th17 –: T helper 17.
TMT –: Tandem mass tag.
Treg –: Regulatory T cell.
UC –: Ulcerative colitis.
WT –: Wild type.

---

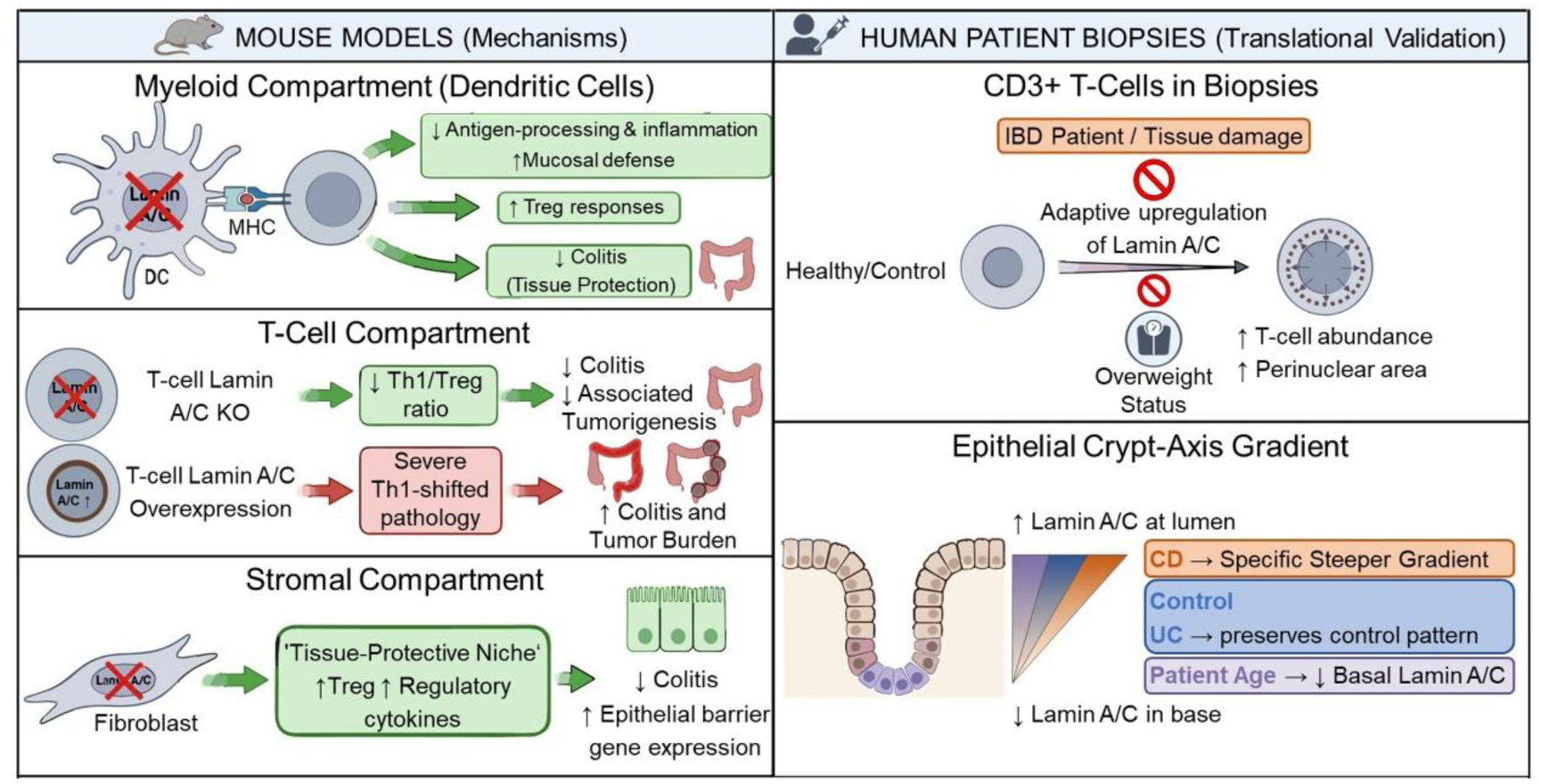
Graphical Abstract. Lamin A/C is a cell-type- and context-dependent regulator of intestinal inflammation, linking nuclear lamina architecture to compartment-specific control of mucosal pathology, serving as a potential therapeutic and biomarker axis in IBD.

**Supplementary Figure 1:**
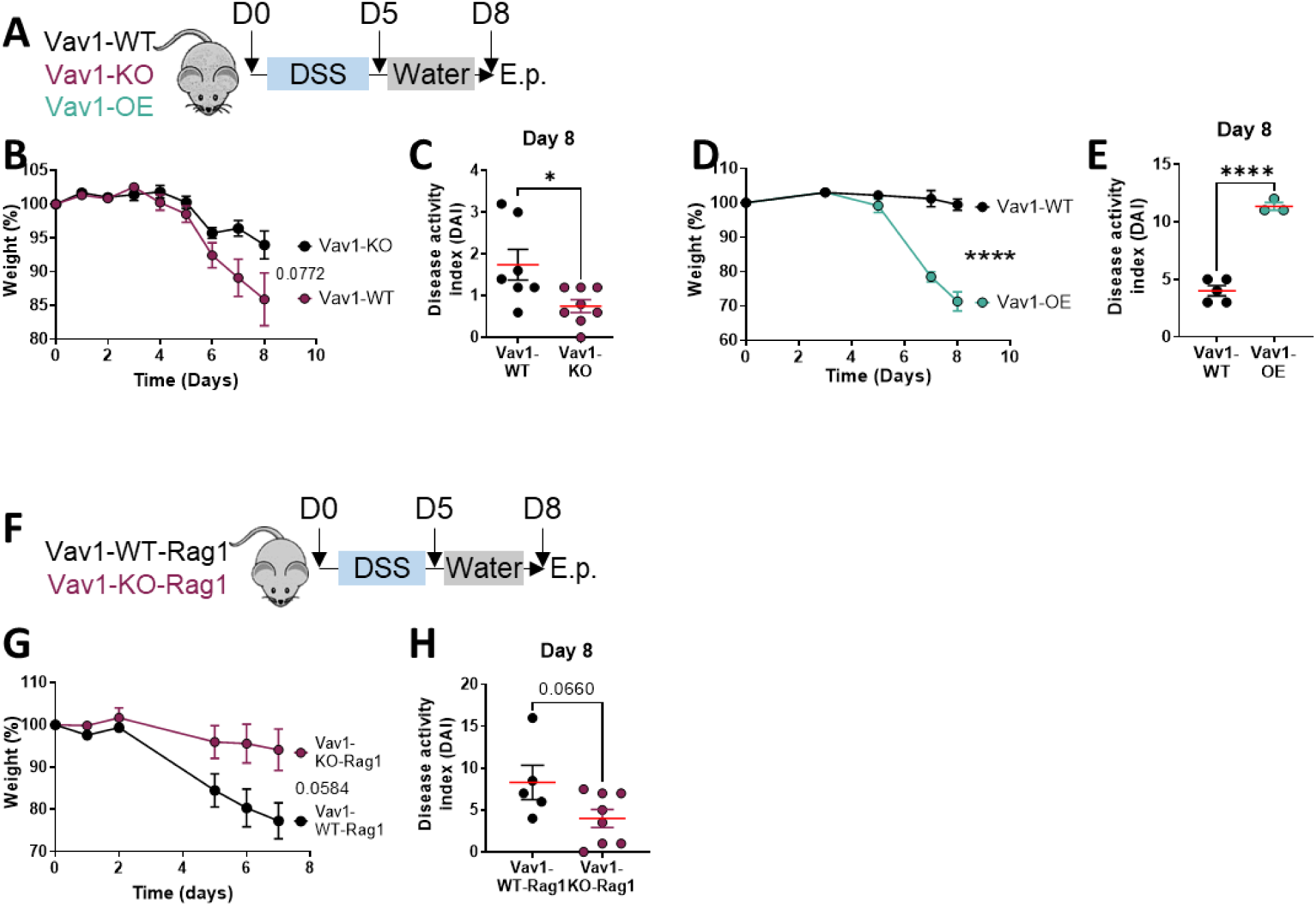
Pan-hematopoietic lamin A/C levels determine susceptibility to acute DSS-induced colitis and confer protection independently of adaptive lymphocytes. (A) Schematic representation of the experimental protocol used to induce acute colitis with a single DSS cycle in wild-type mice (Vav1-WT), pan-hematopoietic lamin A/C-deficient mice (Vav1-KO), and pan-hematopoietic lamin A/C-overexpressing mice (Vav1-OE). (B) Body weight evolution during DSS exposure, expressed as percentage of day 0, in Vav1-WT and Vav1-KO mice. (C) Disease activity index (DAI) at day 8 e.p. in Vav1-WT and Vav1-KO mice. (D) Body weight evolution during DSS exposure, expressed as percentage of day 0, in Vav1-WT and Vav1-OE mice. (E) DAI at day 8 e.p. in Vav1-WT and Vav1-OE mice. (F) Schematic representation of the acute DSS colitis protocol in Vav1-WT-Rag1 and Vav1-KO-Rag1 mice, both lacking mature T and B cells due to Rag1 deficiency, with Vav1-WT-Rag1 mice maintaining physiological lamin A/C levels and Vav1-KO-Rag1 mice lacking lamin A/C in the pan-hematopoietic compartment. (G) Body weight evolution during DSS exposure, expressed as percentage of day 0, in Vav1-WT-Rag1 and Vav1-KO-Rag1 mice. (H) DAI at day 8 e.p. in Vav1-WT-Rag1 and Vav1-KO-Rag1 mice. Data are shown as mean ± SEM (n = 3–8 mice per genotype for Vav1-WT/Vav1-KO/Vav1-OE comparisons; n = 5–8 mice per genotype for Vav1-WT-Rag1 and Vav1-KO-Rag1). Statistical significance was assessed using two-way ANOVA with Tukey’s multiple-comparison test for longitudinal measurements and unpaired Student’s t-test for endpoint analyses; *p < 0.05, **p < 0.01, ***p < 0.001, ****p < 0.0001; trends (p < 0.1) are indicated where applicable. Endpoint: e.p.

**Supplementary Figure 2:**
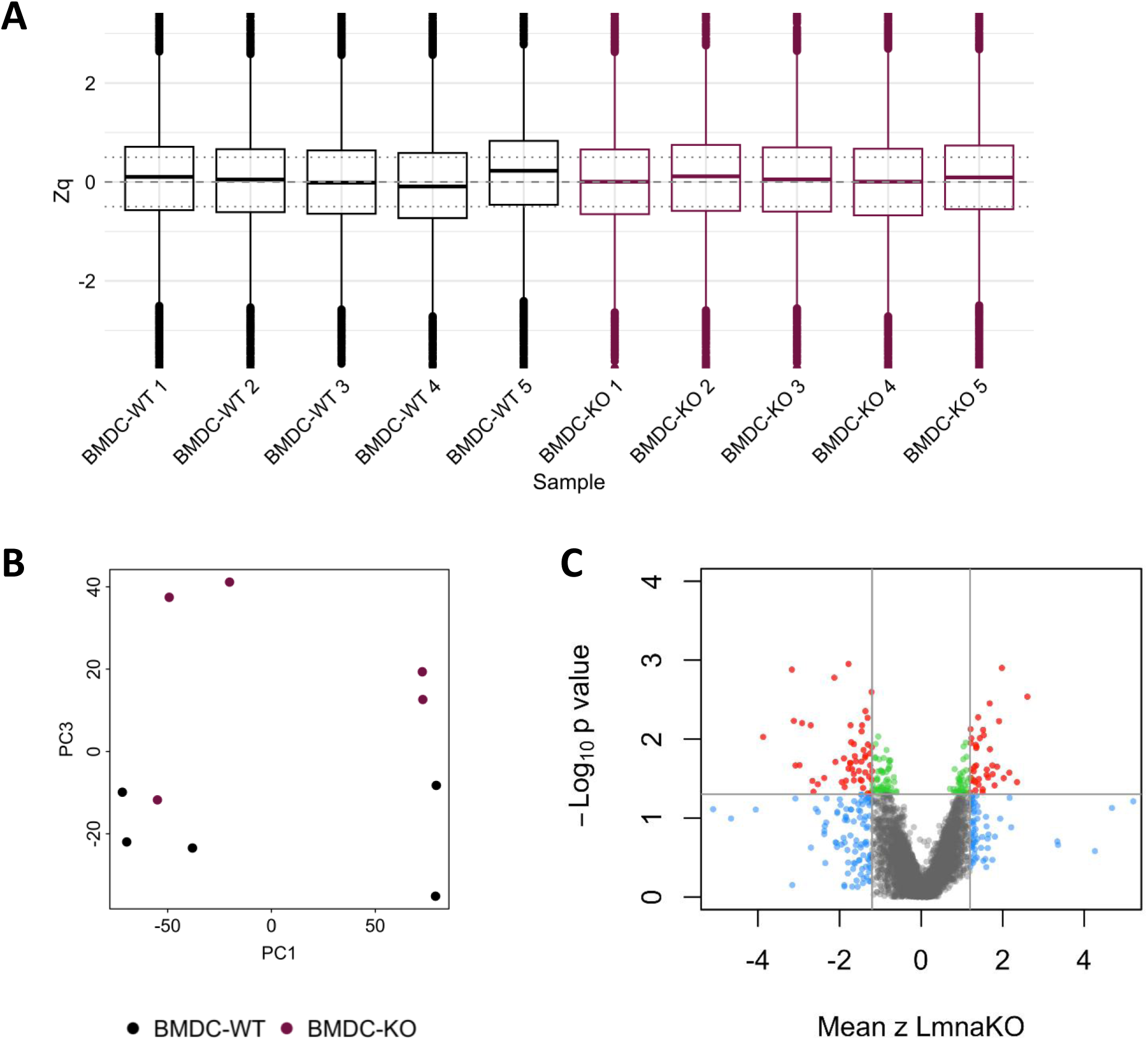
GM-CSF-generated-LPS-matured–lamin A/C–deficient bone marrow-derived dendritic cells (BMDC) proteomics exploration analyses: (A) Boxplot to assess whether samples were correctly normalized. All the samples show their mean (thick line) within the normality threshold area, delimited by the dotted lines (±0.5). (B) Principal component (PC) analysis to assess within-group and between-group variability. Group separation is shown by the third component (cumulative proportion of variance explained 65.8). (C) Volcano plot for statistical significance threshold determination. The x axis displays the mean z-scores from the WSPP model across samples, and the y axis, p values from the limma models. The horizontal dashed line indicates the optimal significance threshold (α= 0.05), and the vertical dashed lines (±1.2) represent the z-score threshold that maximize the proportion of red points (significant proteins with high effect size) in relation to blue points (non-significant with high effect size), green points (significant with low effect size), and grey points (non-significant with low effect size).

**Supplementary Figure 3:**
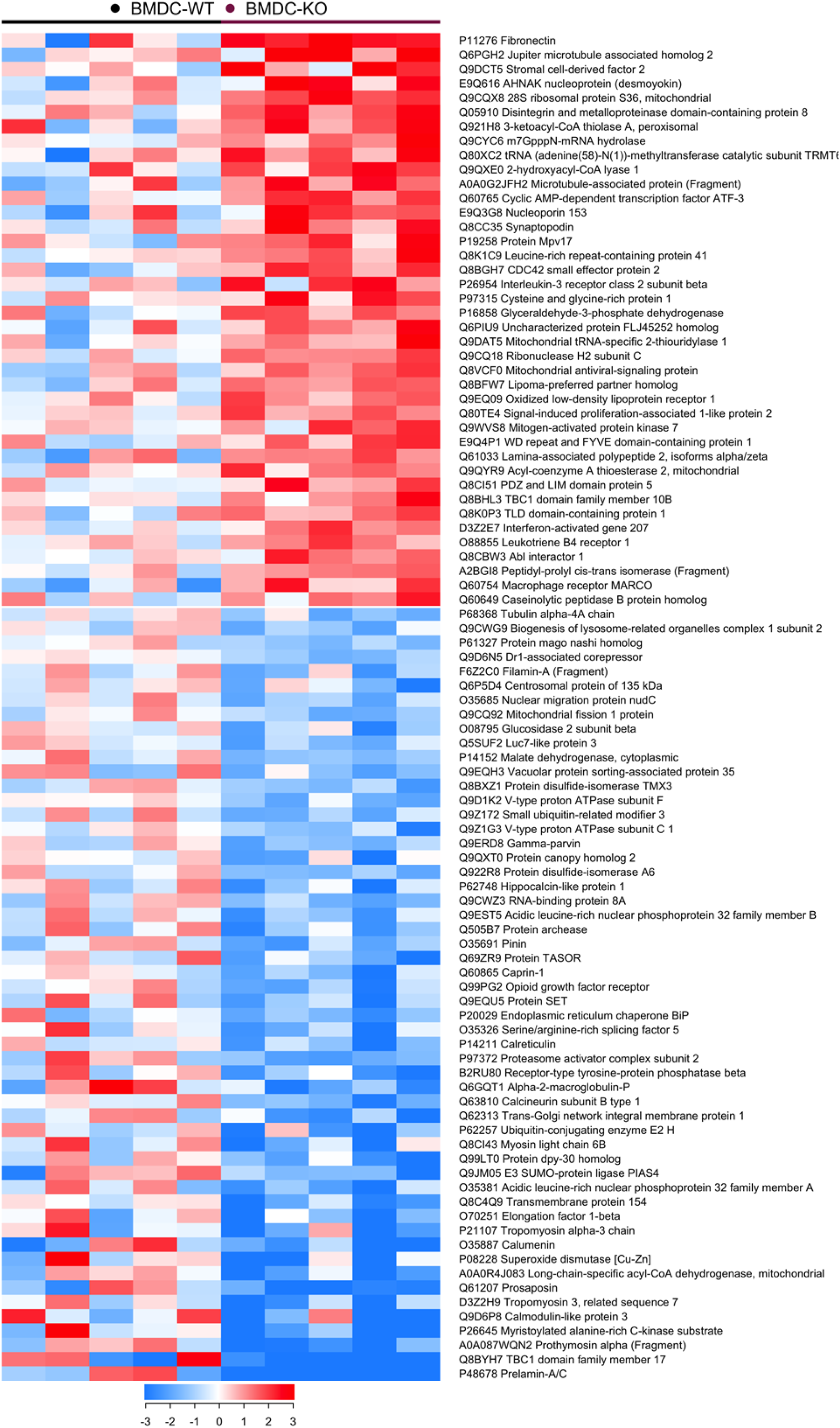
GM-CSF-generated-LPS-matured–lamin A/C–deficient bone marrow-derived dendritic cells (BMDC) proteomics exploration analyses. Relative abundance changes in each sample. Heat map of curated protein candidates with altered expression compared with WT (|mean z| ≥ 1.2 and p value < 0.05). The internal gradient color reflects the individual protein z-scores, with color intensity saturating at |z| ≥ 3. Red: upregulation; blue: downregulation.

**Supplementary Figure 4:**
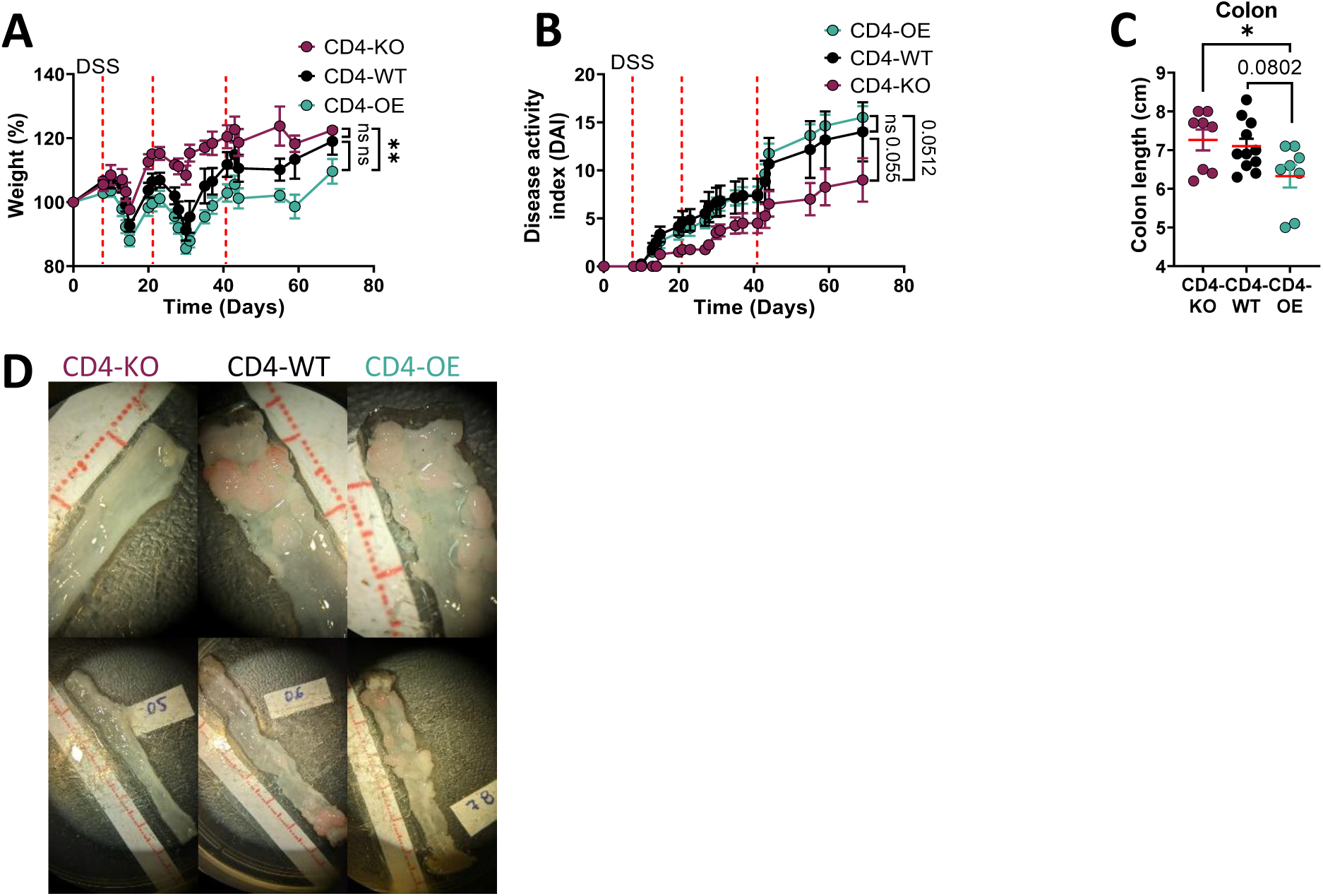
Colitis parameters and colon images in the AOM/DSS colitis-associated colorectal cancer model, A/C in wild-type (CD4-WT), T cell–specific lamin A/C–deficient (CD4-KO), and T cell–specific lamin A/C–overexpressing (CD4-OE) mice. (A) Body weight evolution during the AOM/DSS protocol, expressed as percentage of day 0, in CD4-WT, CD4-KO, and CD4-OE mice. Discontinuous vertical lines indicate the start of each DSS cycle. (B) Disease activity index (DAI) during the AOM/DSS protocol. (C) Colon length measured at sacrifice. (D) Representative images of tumors. Data are shown as mean ± SEM (n = 7–12 mice per genotype pooled from two independent experiments). Statistical significance was assessed using two-way ANOVA with Tukey’s post hoc test for longitudinal measurements and unpaired Student’s t-test for endpoint analyses; *p < 0.05, **p < 0.01; trends (p < 0.1) are indicated where applicable.

**Supplementary Figure 5:**
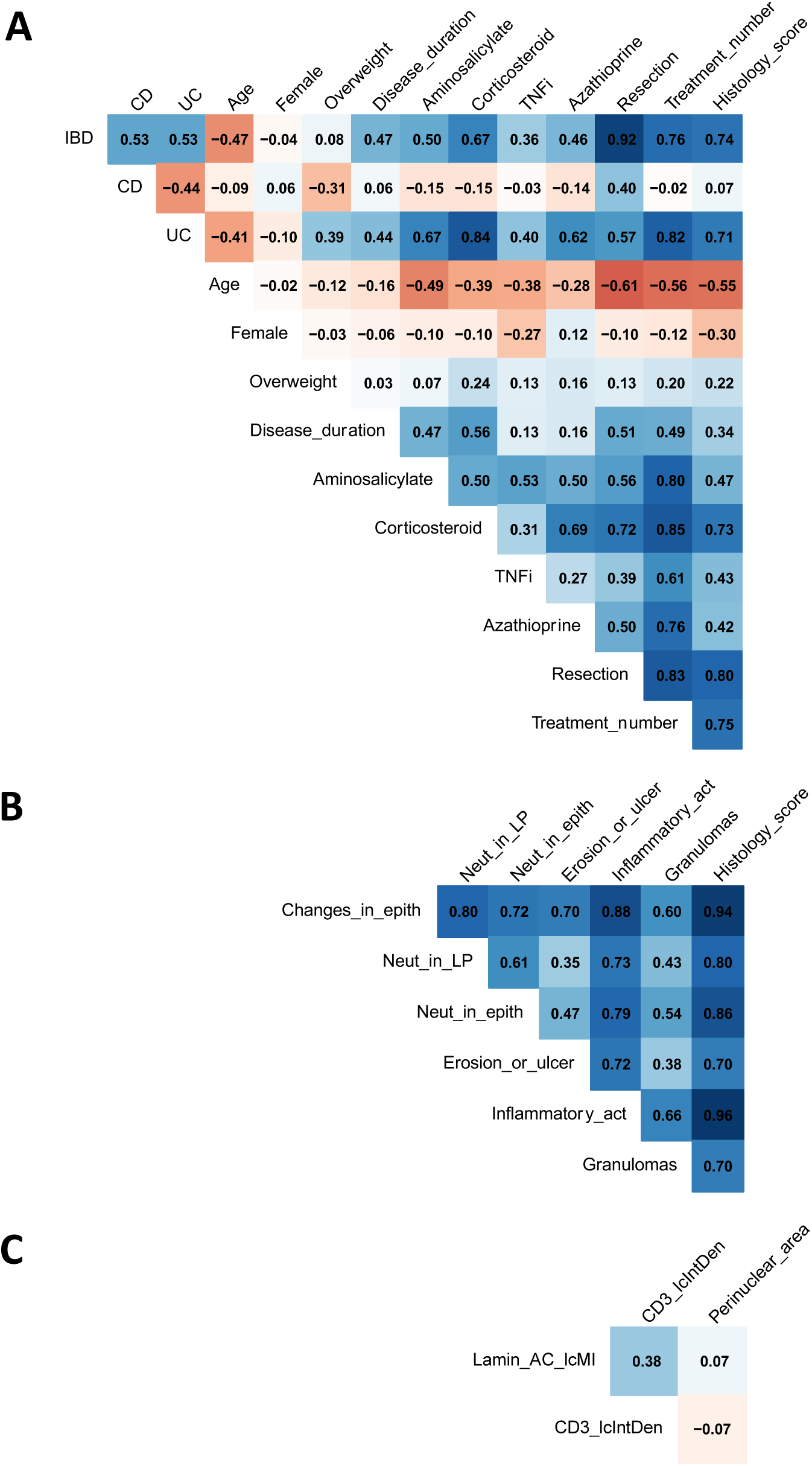
Pearson’s Correlations between Variables of human IBD patients. A) Demographics, clinical characteristics, treatment, and averaged histology score at patient level. B) Histopathological characteristics at ROI level. C) Lamina propria T lymphocyte cellular Characteristics. Crypt characteristics were not represented because there is only one potential covariate.

**Supplementary Figure 6:**
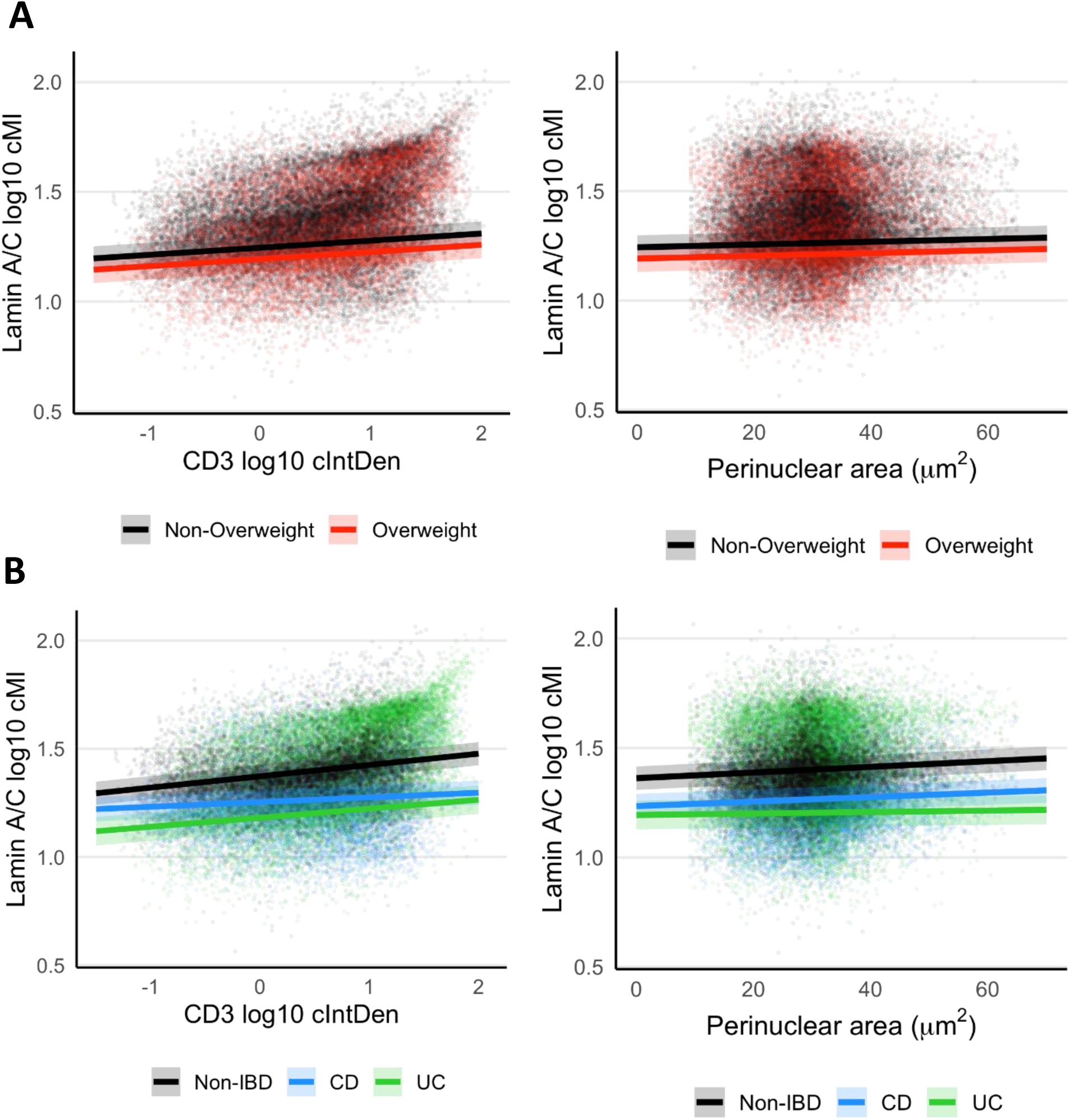
Lamin A/C Expression in CD3+ Cells in Human IBD. Lamin A/C corrected mean intensity (cMI) vs CD3 corrected integrated density (cIntDen) or perinuclear area by A) Overweight status (IBD status main model), B) Group (IBD subtype secondary model).

**Supplementary Figure 7:**
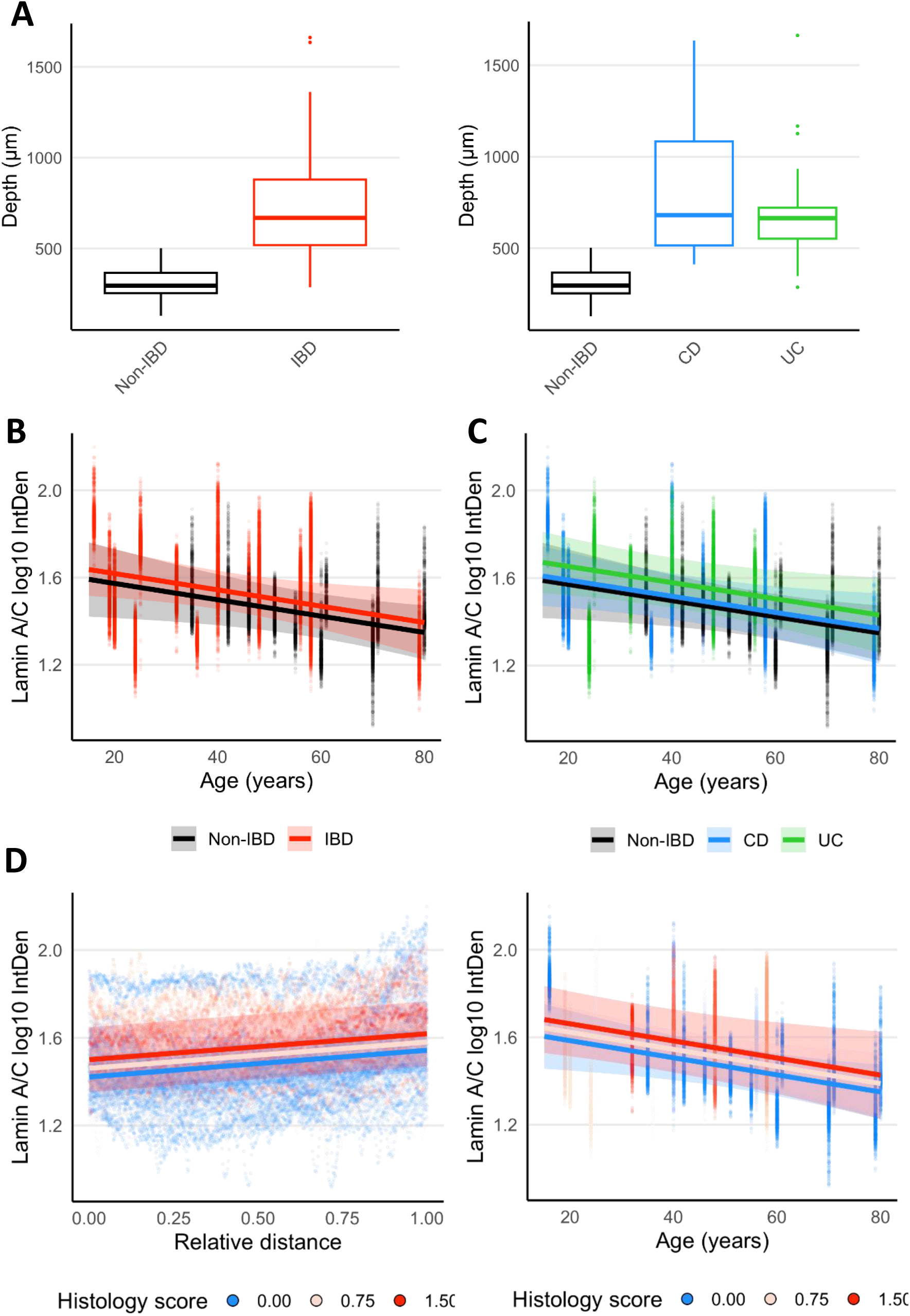
Depth and Lamin A/C Expression in Intestinal Crypts in Human IBD. A) Crypts were significantly deeper in IBD tissues compared to non-IBD controls (p < 0.001), with pronounced hyperplasia observed in both CD (p < 0.001) and UC (p < 0.001). Statistics derived from linear mixed-effects models adjusting for nested random effects (batch, patient). B) Lamin A/C integrated density (IntDen) vs age by IBD status (main model). C) Lamin A/C IntDen vs age by IBD subtype (secondary model). D) Lamin A/C IntDen vs relative distance or age by histology status (secondary model).

**Supplementary Table 1:**
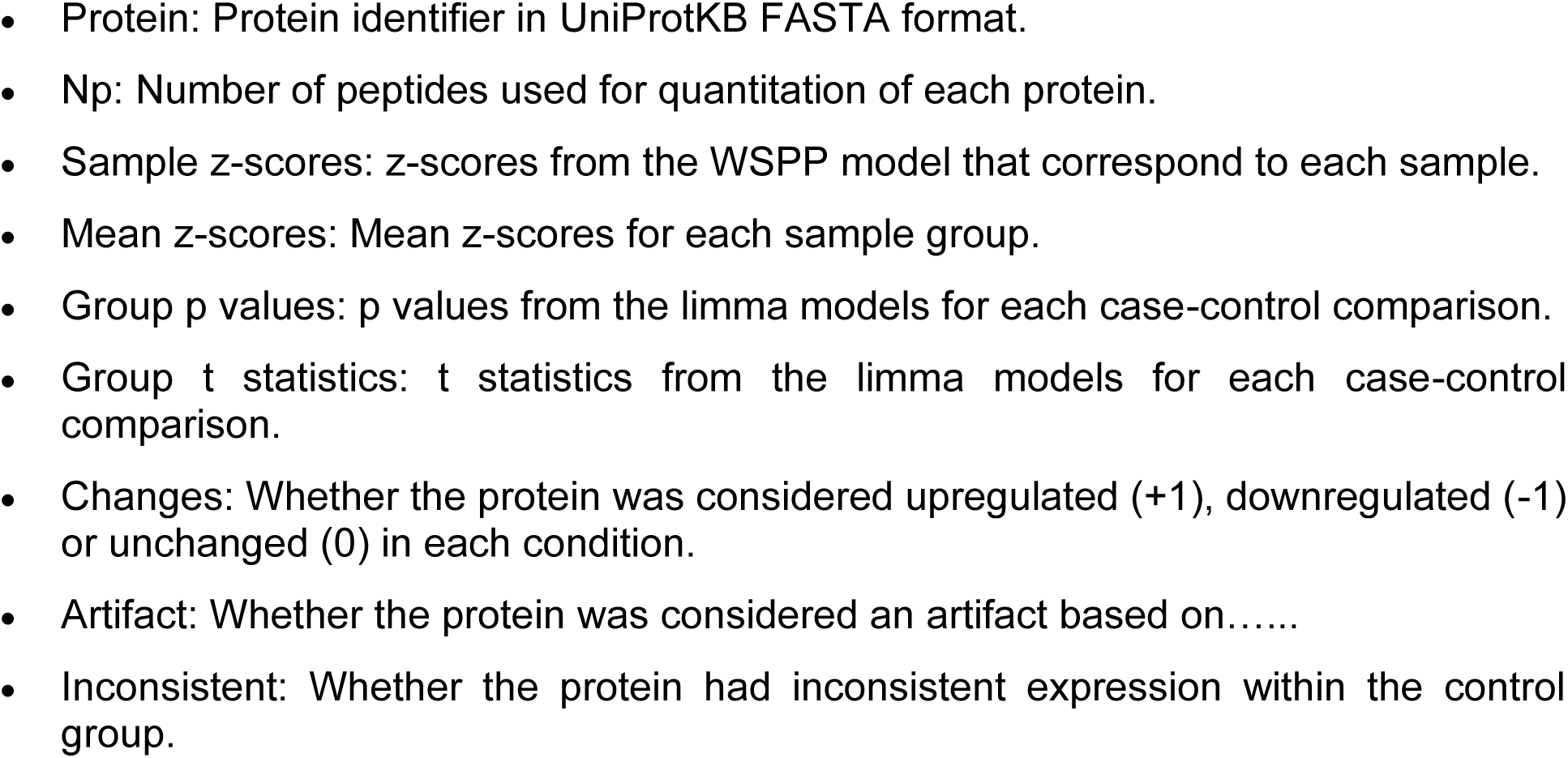
Protein results provided as a separate file, which contains:

**Supplementary Table 2:** Descriptive Analysis of Human IBD Patient Characteristics.

| n (%) | Non-IBD<br>(N=10) | IBD (N=16) |  |  | Overall<br>(N=26) |
| --- | --- | --- | --- | --- | --- |
|  |  | CD (N=8) | UC (N=8) | Overall (N=16) |  |
| Demographics & Clinical Characteristics |  |  |  |  |  |
| CD | 0 (0.0) | 8 (100.0) | 0 (0.0) | 8 (50.0) | 8 (30.8) |
| UC | 0 (0.0) | 0 (0.0) | 8 (100.0) | 8 (50.0) | 8 (30.8) |
| Age, y* | 57.5 (47.3, 67.8) | 40.0 (28.0, 58.0) | 36.0 (24.8, 48.0) | 40.0 (24.5, 52.0) | 48.0 (35.0, 58.0) |
| Female | 6 (60.0) | 5 (62.5) | 4 (50.0) | 9 (56.3) | 15 (57.7) |
| Overweight | 3 (30.0) | 1 (12.5) | 5 (62.5) | 6 (37.5) | 9 (34.6) |
| Disease duration, y* | 0.0 (0.0, 0.0) | 0.0 (0.0, 7.0) | 6.0 (1.8, 14.8) | 6.0 (0.0, 11.0) | 0.0 (0.0, 6.0) |
| Treatment |  |  |  |  |  |
| Aminosalicylate | 1 (10.0) | 2 (25.0) | 7 (87.5) | 9 (56.3) | 10 (38.5) |
| Corticosteroid | 0 (0.0) | 2 (25.0) | 8 (100.0) | 10 (62.5) | 10 (38.5) |
| TNFi | 0 (0.0) | 1 (12.5) | 3 (37.5) | 4 (25.0) | 4 (15.4) |
| Azathioprine | 0 (0.0) | 1 (12.5) | 5 (62.5) | 6 (37.5) | 6 (23.1) |
| Resection | 0 (0.0) | 7 (87.5) | 8 (100.0) | 15 (93.8) | 15 (57.7) |
| Treatment number* | 0.0 (0.0, 0.0) | 2.0 (1.0, 2.5) | 4.0 (3.8, 4.3) | 3.0 (2.0, 4.0) | 1.0 (0.0, 3.0) |
| Histopathological Characteristics*† |  |  |  |  |  |
| Changes in epithelium | 0.0 (0.0, 0.0) | 1.0 (0.4, 1.0) | 1.8 (1.0, 2.0) | 1.0 (0.9, 1.6) | 0.5 (0.0, 1.0) |
| Neutrophils in lamina propria | 0.0 (0.0, 0.0) | 1.0 (0.9, 1.0) | 1.0 (1.0, 1.0) | 1.0 (1.0, 1.0) | 1.0 (0.0, 1.0) |
| Neutrophils in epithelium | 0.0 (0.0, 0.0) | 0.8 (0.0, 1.0) | 1.0 (0.9, 1.3) | 1.0 (0.4, 1.0) | 0.0 (0.0, 1.0) |
| Erosion or ulcers | 0.0 (0.0, 0.0) | 0.0 (0.0, 0.0) | 0.5 (0.0, 0.6) | 0.0 (0.0, 0.5) | 0.0 (0.0, 0.0) |
| Inflammatory activity | 0.0 (0.0, 0.0) | 1.0 (0.8, 1.0) | 1.5 (1.0, 2.1) | 1.0 (1.0, 1.6) | 0.8 (0.0, 1.0) |
| Granulomas | 0.0 (0.0, 0.0) | 0.0 (0.0, 0.5) | 0.3 (0.0, 1.0) | 0.0 (0.0, 0.5) | 0.0 (0.0, 0.4) |
| Histology score | 0.0 (0.0, 0.0) | 0.6 (0.4, 0.7) | 1.1 (0.6, 1.3) | 0.7 (0.6, 1.0) | 0.5 (0.0, 0.7) |
| Lamina Propria T Lymphocyte Cellular Characteristics*‡ |  |  |  |  |  |
| CD3 log<sub>10</sub> cIntDen | 0.4 (0.3, 0.5) | 0.4 (0.3, 0.5) | 0.7 (0.4, 0.9) | 0.5 (0.4, 0.8) | 0.4 (0.3, 0.6) |
| Perinuclear area, μm<sup>2</sup> | 28.2 (27.6, 29.7) | 30.3 (28.5, 32.8) | 30.6 (30.3, 31.0) | 30.5 (29.5, 31.7) | 30.1 (28.3, 30.7) |
| Lamin A/C log<sub>10</sub> cMI | 1.4 (1.3, 1.4) | 1.3 (1.2, 1.4) | 1.4 (1.3, 1.5) | 1.3 (1.2, 1.5) | 1.3 (1.2, 1.4) |
| Intestinal Crypt Characteristics*§ |  |  |  |  |  |
| Lamin A/C log<sub>10</sub> IntDen | 1.5 (1.4, 1.5) | 1.5 (1.3, 1.6) | 1.6 (1.6, 1.6) | 1.6 (1.4, 1.6) | 1.5 (1.4, 1.6) |
| Depth, μm | 288 (278, 312) | 657 (571, 951) | 704 (605, 873) | 680 (574, 915) | 567 (301, 816) |

\* Variables summarized as median (Q1, Q3) instead of n (%) because of their numerical nature. † Information obtained at region of interest level and averaged for each patient. ‡ Information obtained at cellular level and averaged for each selected region, each region of interest and each patient in a nested manner. § Information obtained at cellular level and averaged for each selected crypt and each patient in a nested manner. CD: Chron's disease, CD3: cluster of differentiation 3, cIntDen: corrected integrated density, cMI: corrected mean intensity, IBD: inflammatory bowel disease, IntDen: integrated density, TNFi: tumor necrosis factor alpha inhibitor, UC: ulcerative colitis.

**Supplementary Table 3:** Stepwise Forward Model Selection for Lamina Propria T Lymphocyte of Human IBD Patients Lamin A/C log_10_ Corrected Mean Intensity.

| Fixed effects | VIF | AIC | BIC | P value |
| --- | --- | --- | --- | --- |
| (No covariates) | - | -73411.7 | -73360.2 | - |
| CD3 IcIntDen | 0.0 | -75534.2 | -75474.1 | <0.001 |
| CD3 IcIntDen + Perinuclear area | 1.0 | -75772.6 | -75704.0 | <0.001 |
| CD3 IcIntDen + Perinuclear area + CD3 min | 1.2 | -75813.4 | -75736.2 | <0.001 |
| CD3 IcIntDen + Perinuclear area + CD3 min + IBD | 1.5 | -75832.5 | -75746.7 | <0.001 |
| CD3 IcIntDen:IBD + Perinuclear area + CD3 min | 1.3 | -75954.8 | -75860.3 | <0.001 |
| CD3 IcIntDen:IBD + Perinuclear area:IBD + CD3 min | 1.6 | -75984.4 | -75881.4 | <0.001 |
| CD3 IcIntDen:IBD + Perinuclear area:IBD + CD3 min + Age | 1.6 | -75982.9 | -75871.3 | 0.507 |
| CD3 IcIntDen:IBD + Perinuclear area:IBD + CD3 min + Female | 1.6 | -75982.8 | -75871.2 | 0.555 |
| **CD3 IcIntDen:IBD + Perinuclear area:IBD + CD3 min + Overweight** | **1.6** | **-75987.0** | **-75875.4** | **0.033** |
| CD3 IcIntDen:IBD + Perinuclear area:IBD + CD3 min + Overweight + Disease duration | 1.6 | -75985.1 | -75865.0 | 0.714 |
| CD3 IcIntDen:IBD + Perinuclear area:IBD + CD3 min + Overweight + Aminosaliclylate | 1.6 | -75985.9 | -75865.7 | 0.350 |
| CD3 IcIntDen:IBD + Perinuclear area:IBD + CD3 min + Overweight + Corticosteroid | 1.6 | -75988.2 | -75868.0 | 0.073 |
| CD3 IcIntDen:IBD + Perinuclear area:IBD + CD3 min + Overweight + TNFi | 1.6 | -75988.6 | -75868.4 | 0.057 |
| CD3 IcIntDen:IBD + Perinuclear area:IBD + CD3 min + Overweight + Azathioprine | 1.6 | -75985.3 | -75865.1 | 0.570 |
| CD3 IcIntDen:IBD + Perinuclear area:IBD + CD3 min + Overweight + Resection | 2.5 | -75985.5 | -75865.4 | 0.455 |
| CD3 IcIntDen:IBD + Perinuclear area:IBD + CD3 min + Overweight + Treatment N | 1.6 | -75985.2 | -75865.1 | 0.625 |
| CD3 IcIntDen:IBD + Perinuclear area:IBD + CD3 min + Overweight + Histology score | 1.6 | -75985.1 | -75864.9 | 0.781 |

All models include nested random effects for region, ROI, patient, and batch. The line marked in bold represents the optimal model. “:” marks interaction terms. AIC: Akaike information criterion, BIC: Bayesian information criterion, CD3: cluster of differentiation 3, IBD: inflammatory bowel disease, IcIntDen: log<sub>10</sub> corrected integrated density, IcMI: log<sub>10</sub> corrected mean intensity, min: minimum intensity threshold, TNFi: tumor necrosis factor alpha inhibitor, VIF: variance inflation factor.

**Supplementary Table 4:** Lamina Propria T Lymphocyte of Human IBD Patients Lamin A/C log_10_ Corrected Mean Intensity Model Coefficients.

| Covariate | Estimate | SE | P value |
| --- | --- | --- | --- |
| **Main model: Lamin A/C log<sub>10</sub> cMI vs IBD status** |  |  |  |
| (Intercept) | 1.082 | 0.036 | <0.001 |
| CD3 log<sub>10</sub> clntDen | 0.052 | 0.001 | <0.001 |
| Perinuclear area | 0.001 | <0.001 | <0.001 |
| CD3 min | 0.005 | <0.001 | <0.001 |
| IBD | -0.122 | 0.027 | <0.001 |
| Overweight | -0.051 | 0.025 | 0.051 |
| CD3 log<sub>10</sub> clntDen:IBD | -0.020 | 0.002 | <0.001 |
| Perinuclear area:IBD | -0.001 | <0.001 | <0.001 |
| **Model: Lamin A/C log<sub>10</sub> cMI vs IBD subtype** |  |  |  |
| (Intercept) | 1.051 | 0.036 | <0.001 |
| CD3 log<sub>10</sub> clntDen | 0.052 | 0.001 | <0.001 |
| CD (vs Non-IBD) | -0.111 | 0.029 | 0.001 |
| UC (vs Non-IBD) | -0.162 | 0.035 | <0.001 |
| Perinuclear area | 0.001 | <0.001 | <0.001 |
| CD3 min | 0.005 | <0.001 | <0.001 |
| CD3 log<sub>10</sub> clntDen:CD | -0.031 | 0.002 | <0.001 |
| CD3 log<sub>10</sub> clntDen:UC | -0.011 | 0.002 | <0.001 |
| Perinuclear area:CD | <0.001 | <0.001 | 0.058 |
| Perinuclear area:UC | -0.001 | <0.001 | <0.001 |
| **Model: Lamin A/C log<sub>10</sub> cMI vs histology score** |  |  |  |
| (Intercept) | 1.053 | 0.042 | <0.001 |
| CD3 log<sub>10</sub> clntDen | 0.049 | 0.001 | <0.001 |
| Histology score | -0.051 | 0.027 | 0.069 |
| Perinuclear area | 0.001 | <0.001 | <0.001 |
| CD3 min | 0.004 | <0.001 | <0.001 |
| CD3 log<sub>10</sub> clntDen:Histology score | -0.019 | 0.002 | <0.001 |
| Perinuclear area:Histology score | -0.001 | <0.001 | <0.001 |

All models include nested random effects for region, ROI, patient, and batch. “:” marks interaction terms. CD: Chron’s disease, CD3: cluster of differentiation 3, clntDen: corrected integrated density, cMI: corrected mean intensity, IBD: inflammatory bowel disease, min: minimum intensity threshold, SE: standard error, UC: ulcerative colitis.

**Supplementary Table 5:** Lamina Propria T Lymphocyte of Human IBD Patients Lamin A/C log_10_ Corrected Mean Intensity Model Random Effects.

| Random effect | Variance (by model) |  |  |
| --- | --- | --- | --- |
|  | IBD status | IBD Subtype | Histology score |
| Sub-region:(ROI:(Patient:Batch)) | 0.005 | 0.005 | 0.004 |
| ROI:(Patient:Batch) | <0.001 | <0.001 | 0.001 |
| Patient:Batch | 0.002 | 0.002 | 0.005 |
| Batch | 0.002 | 0.001 | 0.002 |
| Residual | 0.008 | 0.008 | 0.008 |

IBD: inflammatory bowel disease, ROI: Region of interest.

**Supplementary Table 6:** Stepwise Forward Model Selection for Intestinal Crypt of Human IBD Patients Lamin A/C log10 Integrated Density.

| Fixed effects | VIF | AIC | BIC | P value |
| --- | --- | --- | --- | --- |
| (No covariates) | - | -94938.8 | -94895.0 | - |
| Relative distance | 0.0 | -103080.6 | -103028.1 | <0.001 |
| Relative distance + IBD | 1.0 | -103081.7 | -103020.4 | 0.078 |
| Relative distance:IBD | 1.0 | -103140.1 | -103070.1 | <0.001 |
| **Relative distance:IBD + Age** | **1.1** | **-103142.8** | **-103064.1** | **0.030** |
| Relative distance:IBD + Age + Female | 1.1 | -103140.8 | -103053.3 | 0.954 |
| Relative distance:IBD + Age + Overweight | 1.1 | -103141.2 | -103053.7 | 0.530 |
| Relative distance:IBD + Age + Disease duration | 1.1 | -103141.2 | -103053.6 | 0.564 |
| Relative distance:IBD + Age + Aminosalicylate | 1.2 | -103141.4 | -103053.9 | 0.439 |
| Relative distance:IBD + Age + Corticosteroid | 1.1 | -103140.8 | -103053.3 | 0.889 |
| Relative distance:IBD + Age + TNFi | 1.2 | -103143.4 | -103055.9 | 0.105 |
| Relative distance:IBD + Age + Azathioprine | 1.1 | -103141.4 | -103053.9 | 0.442 |
| Relative distance:IBD + Age + Resection | 3.0 | -103143.6 | -103056.1 | 0.094 |
| Relative distance:IBD + Age + Treatment N | 1.4 | -103142.2 | -103054.7 | 0.245 |
| Relative distance:IBD + Age + Histology score | 1.3 | -103141.0 | -103053.5 | 0.660 |

All models include nested random effects for crypt, patient, and batch. The line marked in bold represents the optimal model. ":" marks interaction terms. AIC: Akaike information criterion, BIC: Bayesian information criterion, IBD: inflammatory bowel disease, TNFi: tumor necrosis factor alpha inhibitor, VIF: variance inflation factor.

**Supplementary Table 7:** Intestinal Crypt of Human IBD Patients Lamin A/C log10 Integrated Density Model Coefficients.

| Covariate | Estimate | SE | P value |
| --- | --- | --- | --- |
| **Main model: Lamin A/C log10 IntDen vs IBD status** |  |  |  |
| (Intercept) | 1.596 | 0.108 | <0.001 |
| Relative distance | 0.103 | 0.003 | <0.001 |
| IBD | 0.034 | 0.059 | 0.567 |
| Age | -0.004 | 0.002 | 0.032 |
| Relative distance:IBD | 0.023 | 0.003 | <0.001 |
| **Model: Lamin A/C log10 IntDen vs IBD subtype** |  |  |  |
| (Intercept) | 1.593 | 0.109 | <0.001 |
| Relative distance | 0.103 | 0.003 | <0.001 |
| CD (vs Non-IBD) | -0.001 | 0.064 | 0.984 |
| UC (vs Non-IBD) | 0.084 | 0.070 | 0.243 |
| Age | -0.004 | 0.002 | 0.035 |
| Relative distance:CD | 0.042 | 0.003 | <0.001 |
| Relative distance:UC | -0.004 | 0.003 | 0.278 |
| **Model: Lamin A/C log10 IntDen vs histology score** |  |  |  |
| (Intercept) | 1.602 | 0.094 | <0.001 |
| Relative distance | 0.122 | 0.002 | <0.001 |
| Histology score | 0.053 | 0.059 | 0.385 |
| Age | -0.004 | 0.002 | 0.018 |
| Relative distance:Histology score | -0.002 | 0.003 | 0.389 |

All models include nested random effects for crypt, patient, and batch. “.” marks interaction terms. CD: Chron’s disease, IntDen: integrated density, IBD: inflammatory bowel disease, SE: standard error, UC: ulcerative colitis.

**Supplementary Table 8:**
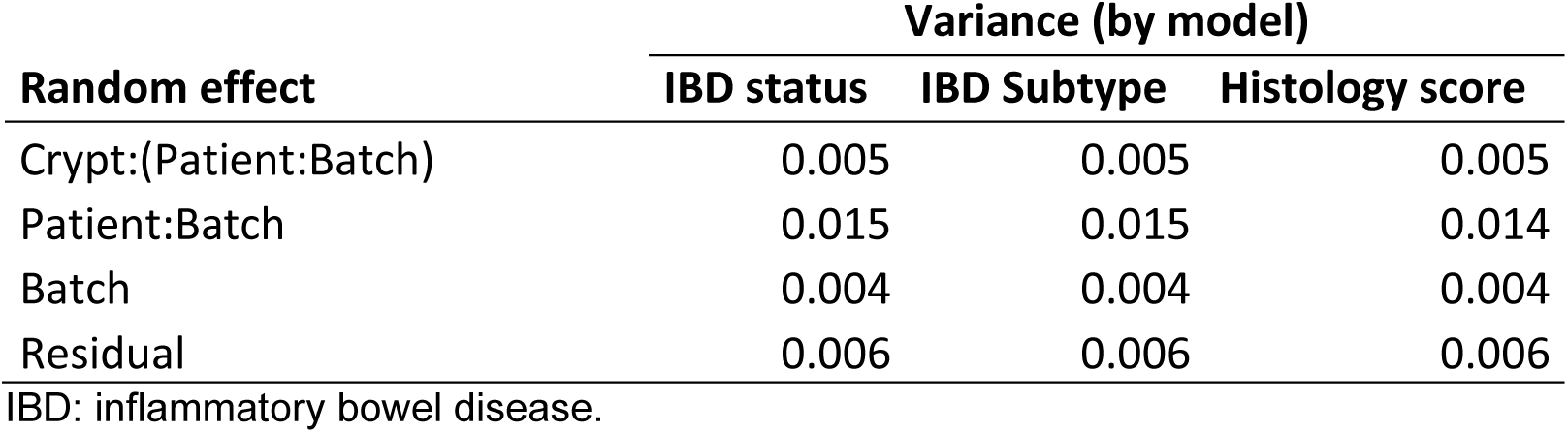
Intestinal Crypt of Human IBD Patients Lamin A/C log10 Integrated Density Model Random Effects.

**Supplementary Table 9:** Flow cytometry fluorophore-conjugated antibodies.

| **Antibody** | **Clone** | **Conjugate** | **Manufacturer** | **Dilution** |
| --- | --- | --- | --- | --- |
| CD3 | 17A2 | APC<br>BV421 | Tonbo<br>BD Horizon | 1:200 |
| CD4 | RM4-5 | V450, APC, PE | Tonbo | 1:200 |
| CD8 | 53-6.7 | PE-Cy5, PE-Cy7 | Tonbo | 1:200 |
| CD25 | PC61.5 | PE-Cy7 | Tonbo | 1:200 |
| CD45.1 | A20 | Pe-Cy7, v450,<br>FITC | Tonbo | 1:200 |
| CD45.2 | 104 | v450, redFluor710 | Tonbo | 1:200 |
| IFNγ | XMG1.2 | FITC, APC | Tonbo | 1:200 |
| Foxp3 | 3G3 | FITC | Tonbo | 1:200 |
| IL10 | JES5-16E3 | FITC, PE | BD<br>Pharmingen | 1:100 |
| RORγt | AFKJS-9 | PE | eBioscience | 1:50 |

**Supplementary Table 10:**
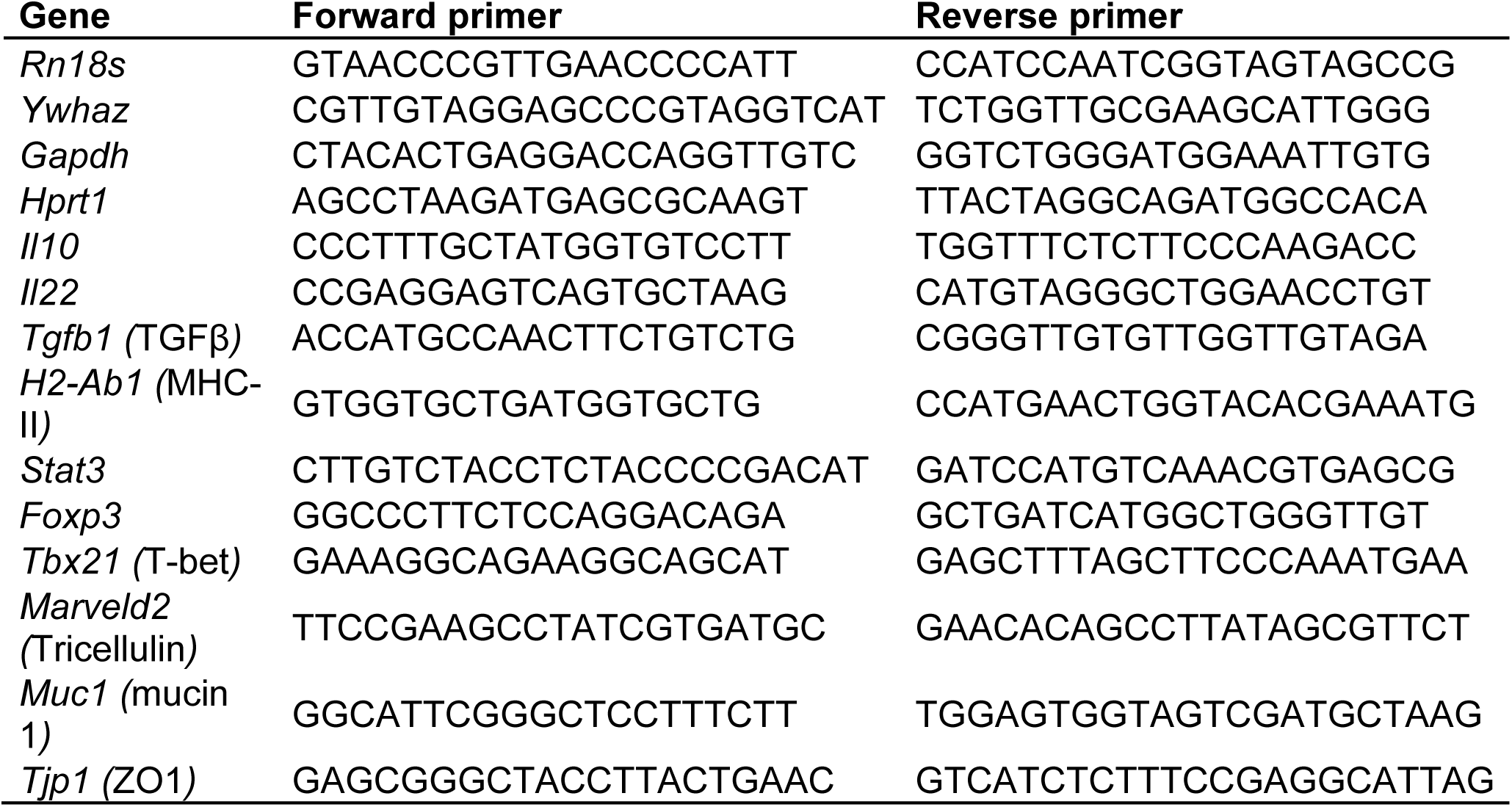
Primer sequences used in q-PCR analysis.

